# A high-content screen profiles cytotoxic microRNAs in pediatric and adult glioblastoma cells and identifies miR-1300 as a potent inducer of cytokinesis failure

**DOI:** 10.1101/789438

**Authors:** Marjorie Boissinot, Henry King, Matthew Adams, Julie Higgins, Thomas A. Ward, Lynette P. Steele, Daniel Tams, Ruth Morton, Euan Polson, Barbara da Silva, Alastair Droop, Josie L. Hayes, Heather Martin, Peter Laslo, Ewan Morrison, Darren C. Tomlinson, Heiko Wurdak, Jacquelyn Bond, Sean E. Lawler, Susan C. Short

**Affiliations:** Radiation Biology and Therapy Group, Leeds Institute of Medical Research, University of Leeds, St James’s Hospital, LS9 7TF, Leeds, UK; Translational Neuro-Oncology Group, Leeds Institute of Medical Research, University of Leeds, St James’s Hospital, LS9 7TF, Leeds, UK; BioScreening Technology Group, University of Leeds, St James’s Hospital, LS9 7TF, Leeds, UK; Stem Cells and Brain Tumour Research Group, Leeds Institute of Medical Research, University of Leeds, St James’s Hospital, LS9 7TF, Leeds, UK; MRC Medical Bioinformatics Centre, University of Leeds, Clarendon Way, Leeds, LS2 9NL, UK; School of Molecular and Cellular Biology, Faculty of Biological Sciences, University of Leeds, LS2 9JT, Leeds, UK; Myeloid Differentiation Group, Leeds Institute of Medical Research, University of Leeds, St James’s Hospital, LS9 7TF, Leeds, UK; Cell Biology Research Group, Leeds Institute of Medical Research, University of Leeds, St James’s Hospital, LS9 7TF, Leeds, UK; Microcephaly and Neurogenesis Research Group, Leeds Institute of Medical Research, University of Leeds, St James’s Hospital, LS9 7TF, Leeds, UK; Harvey Cushing Neurooncology Laboratories, Dept of Neurosurgery, Brigham and Women’s Hospital, Harvard Medical School, Boston, MA02115, USA; St James’s Institute of Oncology and Leeds Institute of Medical Research, University of Leeds, St James’s Hospital, LS9 7TF, Leeds, UK

**Keywords:** MicroRNAs, Glioblastoma, miR-1300, cytokinesis failure, ECT2

## Abstract

**Background:** MicroRNAs play an important role in the regulation of mRNA translation, and have therapeutic potential in cancer and other diseases.

**Methods:** To profile the landscape of microRNAs with significant cytotoxicity in the context of glioblastoma (GBM), we performed a high-throughput screen using a synthetic oligonucleotide library representing all known human microRNAs in adult and pediatric GBM cells. Bio-informatics analysis were used to refine this list and the top seven microRNAs were validated in a larger panel of cells by flow-cytometry, and RTqPCR. The downstream mechanism of the strongest and most consistent candidate was investigated by siRNAs, 3’UTR luciferase assays and Western Blotting.

**Results:** Our screen identified ∼100 significantly cytotoxic microRNAs with 70% concordance between cell lines. MicroRNA-1300 (miR-1300) was the most potent and robust candidate. We observed a striking binucleated phenotype in miR-1300 expressing cells and characterized the mechanism of action as cytokinesis failure followed by apoptosis, which was observed in an extended GBM cell panel including two stem-like patient-derived cultures. We identified the physiological role of miR-1300 as a regulator of endomitosis in megakaryocyte differentiation where blockade of cytokinesis is an essential step. In glioblastoma cells, the oncogene Epithelial Cell Transforming 2 (ECT2) was validated as a direct key target of miR-1300. ECT2 siRNA phenocopied the effects of miR-1300, and its overexpression led to a significant rescue of miR-1300 induced binucleation.

**Conclusion:** MiR-1300 was identified as a novel regulator of endomitosis with translatable potential for therapeutic application. The datatasets will be a resource for the neuro-oncology community.

**Key points (2 or 3 key points 85 characters plus spaces each):** 70% of cytotoxic microRNAs were shared between adult and pediatric glioblastoma cells

MiR-1300 expression is restricted to endomitosis within megakaryocyte differentiation

MiR-1300’s ectopic expression is a potent and promising therapeutic tool in cancer

**Importance of Study:** Previous functional studies of microRNAs involved in the regulation of glioblastoma cell proliferation and/or survival have focused on adult glioblastoma alone and are restricted to only a few microRNAs at a time. Our study provides the first encompassing landscape of potent cytotoxic microRNAs in pediatric and adult glioblastoma.

Not only, does our data provide an invaluable resource for the research community but it also revealed that 70% of microRNAs with significant cytotoxicity were shared by adult and pediatric cells. Finally, we identified and characterized the previously undescribed role of microRNA-1300 in the tight regulation of megakaryocyte differentiation into platelets and how, when expressed outside of this context, miR-1300 consistently causes cytokinesis failure followed by apoptosis, and thus represents a powerful cytotoxic tool with potential for translation towards therapeutic applications.

## Introduction

MicroRNAs are small 22-24nt single-stranded non-coding RNAs that function by reducing the translation of target mRNAs. In glioblastoma (GBM), they have been shown to play roles in proliferation, invasion and stemness, suggesting that microRNAs and their downstream pathways may represent potent therapeutic targets (1-6). There is an increasing number of microRNA mimics and inhibitors in pre-clinical and early clinical development in cancer, including one for patients with solid tumors using a mimic of microRNA-34 (MRX34, Mirna Therapeutics Inc., NCT01829971) (7-10). Recent pre-clinical studies showed efficacy of a microRNA expressing therapeutic vector in GBM (11). MicroRNA-10b expression has been measured in a clinical trial (NCT01849952) to assess its use as a prognostic and diagnostic biomarker. An inhibitor of miR-10b is also currently at the preclinical development stage (Regulus Therapeutics Inc. and (12)).

Current approaches for microRNA studies in GBM mainly involve endogenous microRNA expression profiles coupled with bioinformatic analysis and target identification to link the landscape of microRNA expression to GBM biology and disease outcome (13-15). Other functional studies have focused on small numbers of microRNAs and very few large scale functional studies have been performed in GBM (16).

To access the landscape of potential cytotoxic microRNAs in GBM we decided on a global approach by performing a large-scale functional screen. We used a microRNA mimic oligonucleotide library combined with a high-throughput imaging platform to identify microRNAs that significantly impaired proliferation and/or survival of GBM cells. This approach highlighted microRNA-1300 as a candidate for more detailed characterization. We found that ectopic expression of the mature form of miR-1300 consistently caused a G2/M cell cycle arrest followed by apoptosis. Further validation showed that miR-1300 caused cytokinesis failure and the oncogene Epithelial Cell Transforming 2 (ECT2) was identified as one of the direct targets of miR-1300 involved in this phenotype. This, in turn, led us to identify the key physiological need for the finely tuned expression of miR-1300 during endomitosis in platelet formation from megakaryocytes (17-19). Taken together, our study not only provides an encompassing profile of cytotoxic microRNAs towards adult and pediatric GBM cells but also identifies miR-1300 as a uniquely specific tool with a potential therapeutic window for combination with current standard therapy in GBM. Our dataset will provide a useful resource for other researchers in the field with an interest in the therapeutic application of microRNAs.

## Material and Methods

### Please see supplementary information for: cell lines details, bioinformatics analysis, immunofluorescence, flow-cytometry, qRTPCR, siRNA Knock-Down, Westernblotting, Cell manipulations

#### High-Throughput Screen

The miRIDIAN microRNA MIMIC library based on miRBase v16.0 was purchased from Dharmacon (GE Healthcare). All microRNA mimics and siRNAs were resuspended at a stock concentration of 20μM in 1X siRNA Buffer prepared from a 5X stock (Cat# B-002000-UB-100, Dharmacon GE Healthcare) in RNase free water (Cat# B-003000-WB-100, Dharmacon GE Healthcare). The screen controls for cell number were MIMIC negative control #1 tagged with Dy547 (Cat# CP-004500-01-20) to calculate transfection efficiency as well as without Dy547 (Cat.# CN-001000-01-20) which were included in 8 separate wells and PLK-1 siRNA SMARTpool (Cat.# M-003290-01-005) which was the positive control for decreased cell number. Reverse transfection was performed on U251 and KNS42 cells using RNAiMAX lipofectamine (Invitrogen, Life Technologies, Cat# 13778-075), OPTI-MEM® I Reduced Serum Medium (Cat# 31985, Gibco™, Invitrogen corporation) and 100nM of RNA material (MIMIC or siRNA candidates and controls). For the screen, each well of a 96 well ViewPlate (6005182, Perkin Elmer), contained 20μl of transfection mix including: 0.1μl of RNAiMAX (137789-075, Life Technologies) commercial stock solution, 0.1μl of 20μM microRNA, and 19.8μl of OptiMEM® media (31985-047, Life Technologies); to which 80μl of growth media, containing 6 ×10^3 cells/well (U251) or 9 × 10^3 cells/well (KNS42), were added giving a final volume of 100μl per well. Plates were placed at 37°C, 5% CO_2_ for 72h prior to being fixed in 4% PFA and stained with DAPI (Cat□ D1306, Life Technologies) for nuclei count, TOTO-3 (Cat□ T-3604, Molecular Probes) and Phalloidin-A488 (Cat□ A12379, Life Technologies) for delimitation of the cytoplasm and actin structure (See the “Immunofluorescence” section below for details).

Each 96 well plate contained one microRNA mimic or microRNA control per well. Plates were imaged on a Perkin-Elmer Operetta High Content Imaging System using Harmony 3.1 software, 3 fluorescent channels, 10 fields/well with a 20x objective. For each cell line, the screen was performed in two separate passages of cells.

An algorithm for nuclei counting based on DAPI stain was designed in Columbus 2.4 analysis software. For the analysis, the mean cell number of 8 mimic negative control wells was evaluated per plate and used to calculate the Z-score (the number of standard deviations above or below the mean cell number) for each mimic microRNA utilized. Candidate microRNAs were identified as those which significantly decreased cell number due to reduced proliferation and/or cell survival only if their mean z-score was greater or equal to two standard deviations below the mean z-score of the negative controls for both biological replicates.

#### Induction of Endomitosis

Endomitosis was induced by culturing non-adherent CMK cells for up to 72h in presence of 5μM of the Src kinase inhibitor SU6656 (Sigma Aldrich) (20, 21). Six milliliters of cell suspension were harvested every day for three consecutive days. 2ml were used to prepare RNA (SU6656 experiments only), 2ml to prepare protein lysates for ECT2 western blot, and 2ml were used for imaging.

#### Forced differentiation of Glioma stem-cells

Glioma stem cells were differentiated in the presence of 100ng/ml BMP4 (22-24) (Life Technologies) for 4-5 days prior to transfection with mimic miRNA-1300 or scrambled control. Cells were maintained in BMP4 supplemented media following transfection.

### 3’UTR target validation assays

The 3’UTR reporter assay was performed using Luc-Pair miR Luciferase Assay Kit (Genecopoeia) as per manufacturer’s instructions. Briefly, 9000 cells were seeded per well into a white 96 well plate. The cells were reverse transfected with 1 μg miRNA (scramble of miR1300) and 1 μg reporter plasmid (Control vector CmiT000001-MT01, *ECT2* 3’UTR-WT wild type HmiT064169-MT01, *ECT2* 3’UTR-mt mutant containing a custom two base pair “mutation” turning the human miR-1300 binding site into the *Mus musculus* miR-1300 seed sequence CS-HmiT064169-MT01-01, Genecopoeia). At 72 h post transfection (120 h in GBM4) the media was removed and luminescence was measured on a Borthold Mithras LB 940 plate reader.

### ECT2 rescue

Reverse transfection of KNS42 cells (9000 cells per well in a 96 well Perkin Elmer ViewPlate, Cat# 6005182) with miR-1300 or scrambled control was performed as outlined above (High throughput screen). After 36 hours incubation cells were transfected with 1 μg of either control vector or a vector containing *ECT2* lacking the 3’UTR region (ECT2-Δ3’UTR) using lipofectamine (EX-NEG-Lv105 and EX-T7673 Lv105 respectively, both from Genecopoeia). At 72h post miRNA transfection the cells were fixed with 4% PFA and stained as outlined in the high-throughput screen. The time point at 36h post transfection with miR-1300 prior to *ECT2* rescue was chosen to allow for the G2/M block to take place without the cells being beyond rescue by being too engaged in apoptosis. In GBM1 cells ECT2 rescue experiments the time course was extended to reflect the difference in growth rate and kinetics of the miR-1300 phenotype; transfection with ECT2-Δ3’UTR took place at ∼60 h and fixing at 120 h.

### Sample size and statistical analysis

With the exception of the high-throughput screen, all assays were performed as three biological replicates (cells from different passage number), each containing three technical replicates (three individual wells receiving the same treatment). In the case of immunofluorescent imaging analysis, a minimum of 100 cells per condition were scored. The unequal variance Welch, unpaired t test was chosen to test how far apart the two populations tested were regardless of the difference in their standard deviation (for example: microRNA-1300 vs control).

## Data Availability

The raw data from the screen are supplied in the supplementary information as individual Excel files.

## Results and Discussion

### A high-throughput screen identifies microRNAs with cytotoxic activity in GBM

We profiled cytotoxic microRNAs in adult and pediatric GBM cells using a high-throughput high-content gain-of-function screen based on a library that encompassed mimics of the mature form of all annotated microRNAs based on miRBase v16.0 at the time. The screen was performed in two established GBM cell lines: U251 (adult GBM) and KNS42 (pediatric GBM). A schematic of the screen is shown in Suppl. Figure 1. The chosen primary end-point read out was a decrease in cell number assessed by automated nuclei counting at 72 hours post-transfection.

The candidate hit list for KNS42 and U251 cells contained 83 and 304 such candidate hits, respectively (see Suppl. Tables 2 and 3 for the complete screen datasets). The U251 list was re-analyzed using a <-3 Standard Deviation cut-off, which gave 111 hits to consider for further analysis. Based on the z-score analysis, we initially observed a 70% overlap between the candidate lists for each cell line (Suppl. Table 1) and focused further investigations on those microRNAs. We first utilized the online microRNA databases miRBase (25-29) and Targetscan (30-36), as well as PubMed (National Center for Biotechnology Information, National Library of Medicine, Bethesda MD, USA) to gather available information for each of the ‘hit’ microRNAs: chromosomal location, main validated and predicted target genes, associated functions and role in disease. This approached allowed us to shortlist 18 candidate microRNAs. The second level of analysis took into consideration the strength of the z-score, together with information focused on seed sequence family, the association of the target genes with cell proliferation and/or cell death, led to the selection of seven of those eighteen microRNAs for validation in cell based assays. We applied the conditions of the primary screen to a panel of four established GBM cell lines (U251, KNS42, LN229 and U373), this confirmed a statistically significant cytotoxicity following transfection of all seven mimic-microRNA candidates (Suppl. Figure 2A).

In order to investigate whether the reduced cell number was due to programmed cell death, we analyzed expression of cleaved caspase-3 by immunofluorescence. Interestingly, only transfection with the mimic for miR-1300 led to the cleavage of caspase-3 and cell death by apoptosis (Suppl. Figure 2B and 2C).

We then focused further on characterizing the role and mechanism of action of miR-1300; its high z-scores (z-score=-2.98 in KNS42 and z-score=-5.14 in U251), and its ability to induce apoptosis making it the most promising and interesting candidate. MiR-1300 mature and precursor sequences as well as alignment to the human genome can be found in supplementary Table 4.

### MiR-1300 induces cytokinesis failure, cell cycle arrest and apoptosis in GBM cell lines and patient-derived GBM

Flow cytometry-based assays were used to measure the effect of ectopic expression of miR-1300 in its mature form (MIMIC) on cell cycle and cell death over time. Expression of miR-1300 induced a significant block of the cell cycle in G2/M at 24 hours in U251 and 48 hours in KNS42 cells (Figure 1A), rapidly followed by the onset of apoptosis (Figure 1B). In U251 cells there was a >60% reduction in cells in G0/G1 with a 1.5-fold increase in G2/M cells at 24 hours compared to the scrambled control (p=0.079**). This G2/M arrest became more pronounced at 48 hours (∼2.5-fold change, p<0.0335*) and 72 hours (3-fold change, p<0.0485*) but was no longer apparent at 96h following high levels of apoptosis (Figure 1C >20% reduction in the number of live cells and ∼3 and 6-fold increase of cells in early-mid and mid-late apoptosis (p=0.0012** and p<0.0001****, respectively). Similar changes in the cell cycle profile were observed in KNS42 cells with a ∼4- and ∼3-fold increase in cells in G2/M at 48 and 72 hours (p=0.017* and p=0.009**, respectively). To further evaluate this effect, we analyzed images of fluorescently stained cells from the screen, which suggested that the cell cycle arrest observed by flow-cytometry occurred during mitosis (Figure 1C). We then formally re-evaluated binucleation in these cells and showed that 72 hours post-transfection, ectopic expression of miR-1300 caused approximately 80% and >60% increase in binucleated cells in U251 and KNS42, respectively (p<0.0001****, in both cases). The same phenotype was observed in LN229 and U373 GBM cell lines (Suppl. Figure 3). The two nuclei per cell which we observed (Figure 1C) suggested that cell cycle arrest took place after telophase and is representative of cytokinesis failure. Live cell imaging confirmed that both KNS42 (Suppl. Movie 1A, link in additional file) and U251 cells (Suppl. Movie 1B, link in additional file) transfected with miR-1300 initiated mitosis normally but failed to complete the final stages of cytokinesis resulting in the formation of binucleated cells.

**Figure 1:**
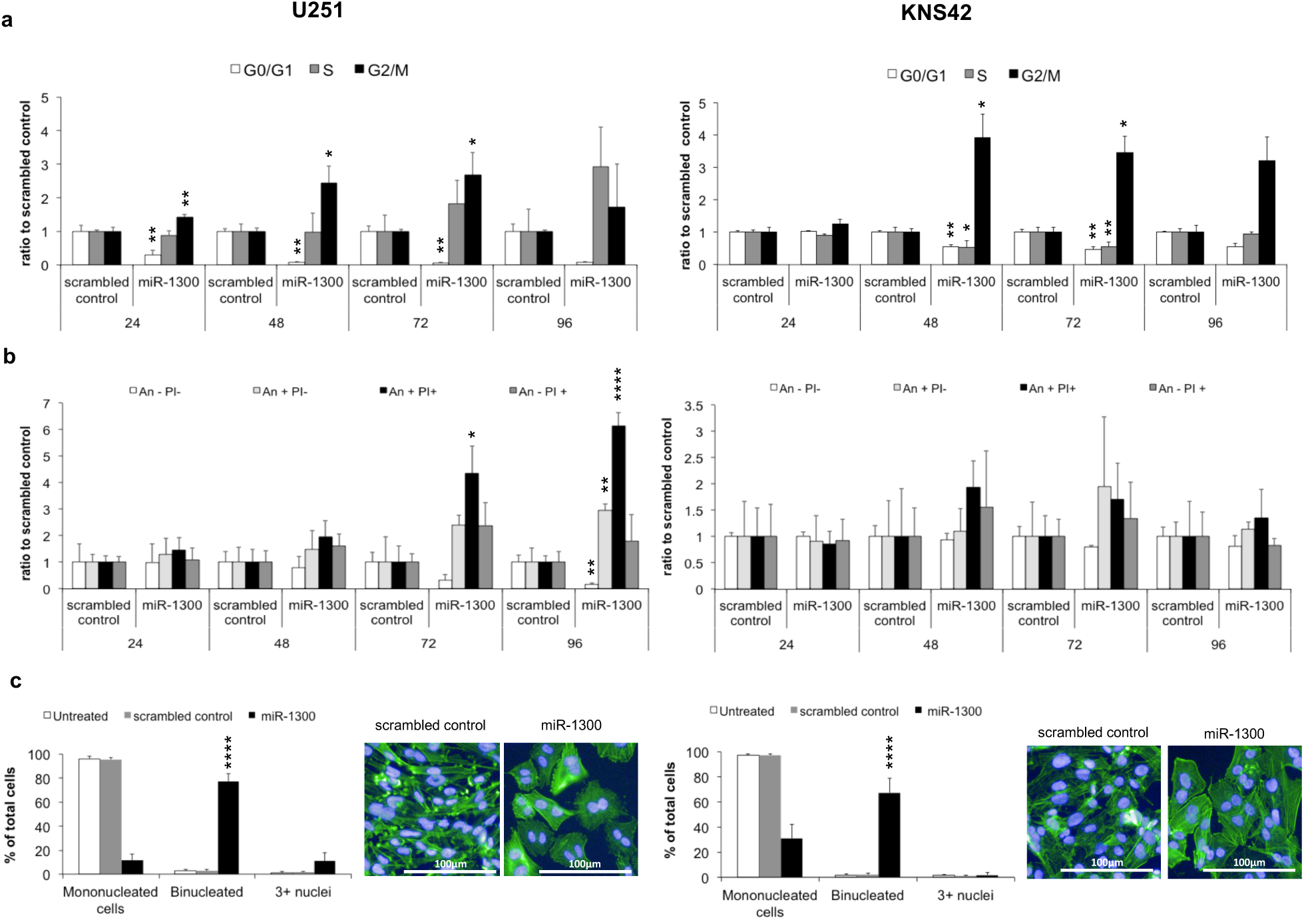
Effect of miR-1300 expression on proliferation and cell death in U251 and KNS42 cell lines **a:** Cell cycle time course analyzed by flow cytometry using propidium iodide (PI) loading as a measure of DNA content. **b:** Cell death measured by flow cytometry where Annexin negative (An -)/ PI negative (PI-) = live cells, An+ PI- = early-mid apoptotic cells, An+ PI+ = mid-late apoptotic cells and An-PI+ = Necrotic cells. **c:** Binucleation phenotype scoring following staining with DAPI and Phalloidin Alexafluor488 (Actin) and the corresponding representative image. All experiments were performed in triplicate. Results are normalized to a scrambled mimic control. Statistical significance is expressed as follows: * = p<0.05, ** = p<0.01, *** = p<0.001 and **** = p<0.0001.

Next, we sought to confirm that this phenotype was also observed in patient-derived glioma stem-like cells (GSCs) as a more representative *in vitro* model. We used two previously characterized GSCs, GBM1 and GBM4 (23, 24) and performed new time courses for the flow cytometry assays at 72, 96 and 120 hours to reflect the slower division rate of the patient-derived cells, and we repeated the imaging experiments with binucleation scoring at 96 hours. Transfection of GBM1 with mimic miR-1300 caused a profound G2/M phase block with a 4-fold increase in the proportion of cells in G2/M phase at 72, 96 and 120 hours compared to cells transfected with the scrambled-mimic control (Figure 2A). Transfection with mimic miR-1300 also caused a 2-fold reduction in the number of cells in G0/G1 and S phase (Figure 2A). This G2/M arrest led to an overall reduction in live cells of approximately 40%, 45% and 80% at 72, 96 and 120 hours and a significant increase (p<0.01 and p<0.05 in GBM1 and GBM4 respectively) in apoptotic cells at 120 hours (Figure 2B). Similar results were observed in GBM4 GSCs with miR-1300 causing a 65% reduction in cells in S-phase at 72 hours which was maintained at 96 hours. At 96 hours there was an increase in cells on G2/M phase by 4-fold for GBM1 and by 2-fold for GBM4 (Figure 2A). At 96 hours, miR-1300 transfection resulted in an approximately 40% reduction in live cells and an increase of around 20% in early-mid apoptotic GBM1 cells. By 120 hours there was an approximately 75% decrease in live cells and a 1.5-fold increase in early-mid apoptotic for both GBM1 and GBM4 and 2.5 and 1.5-fold increase in mid-late apoptotic cells for GBM1 and GBM4 respectively (figure 2B). Figures 2C and D show the effect of miR-1300 on the number of bi- and multi-nucleated cells. As with the established cell lines, miR-1300 caused a decrease in mono-nucleated cells of approximately 90% in both GBM1 and GBM4 and a significant increase (approximately 50%) in bi-nucleated cells for both GSCs, (p=0.0055** and p=0.0081**, respectively).

**Figure 2:**
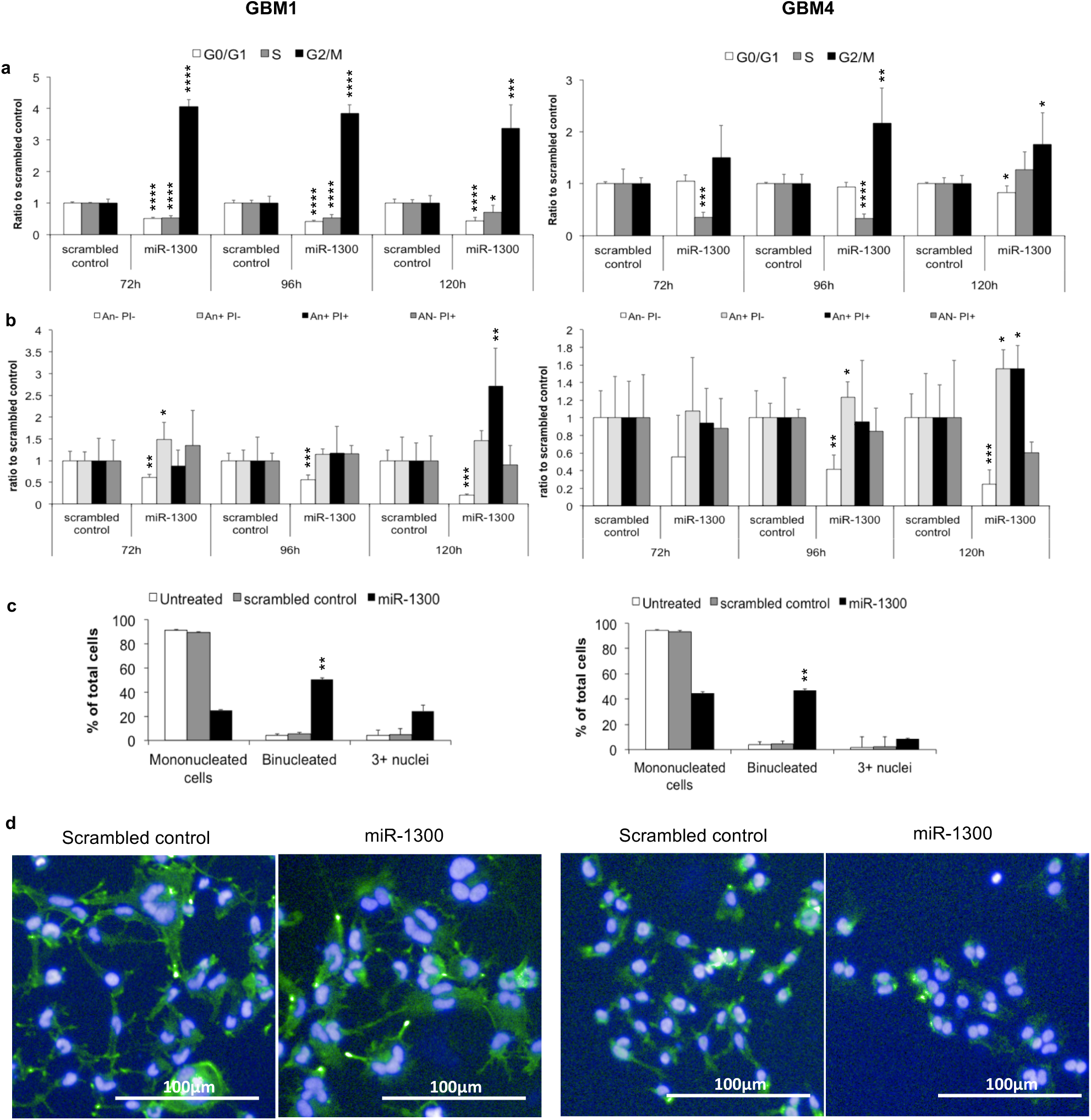
Effect of miR-1300 expression on proliferation and cell death in GSC cultures GBM1 and GBM4. **a**: Cell cycle time course analyzed by flow cytometry using propidium iodide (PI) loading as a measure of DNA content. **b**: Cell death measured by flow cytometry where Annexin negative (An -), PI negative (PI-) = live cells, An+ PI- = early-mid apoptotic cells, An+ PI+ = mid-late apoptotic cells and An-PI+ = Necrotic cells. **c** and **d**: Binucleation phenotype scoring following staining with DAPI and Phalloidin Alexafluor488 (Actin) (3-6 images per condition, representing at least 100 cells) was performed on images taken on the Operetta imaging platform (x10 objective). All experiments were performed in triplicate. Results are normalized to a scrambled mimic control. Statistical significance is expressed as follows: * = p<0.05, ** = p<0.01, *** = p<0.001 and **** = p<0.0001.

Taken together, this initial characterization showed that ectopic expression of miR-1300 consistently led to failure of cytokinesis, measured by cell cycle arrest in G2/M phase and manifested by a striking binucleated phenotype followed by the onset of apoptosis as documented by caspase-3 cleavage (Suppl. Figure 2B/C) and expression of AnnexinV/PI (Figures 1B and 2B). This was confirmed both in established GBM cell lines and in patient-derived GBM cultures.

### Identification of miR-1300 target genes

Since there are no validated target genes for miR-1300, we used the list of predicted targets extracted from the online database TargetScan v5.2 (34). This contains 3327 target genes predicted to be targeted by miR-1300 irrespective of the presence of conserved sites (the seed sequence for miR-1300 is poorly conserved across species). We then loaded this list in the Metacore gene analysis software (Thomson Reuters) and cross-referenced it using AmiGO (37) for “cytokinesis” as a Gene Ontology term. This allowed us to identify which of the miR-1300 target genes had the highest potential involvement in the observed cytokinesis failure. Our analysis gave a list of 21 potential target genes (Suppl. Table 5). Based on the characteristic binucleation seen in our phenotype, the Targetscan prediction score, and the literature, we chose the guanine nucleotide exchange factor (GEF) *ECT2* (*Epithelial Cell Transforming 2*) as our target of interest for initial validation. Interestingly, *ECT2* has been previously described as an oncogene and has been shown to contribute to the invasive behavior of GBM cells (38-42). *ECT2* plays a crucial role in cytokinesis through activation of the small GTPase RhoA, a key protein in the formation of the mitotic cleavage furrow during cytokinesis (17, 18). It has also been shown that treatment of cells with RhoA inhibitors caused a binucleated phenotype similar to that observed in the GBM cell lines following transfection with miR-1300 (43). Further, ECT2 depletion has been shown to lead to cytokinesis failure by impairment of cleavage-furrow formation (44).

Using Real-Time qPCR in a panel of five patient-derived stem-like GBM cell cultures, we observed an inverse relation between *ECT2* mRNA and miR-1300 expression (Suppl. Figure 4A). This is consistent with the frequency of multinuclear cells observed during cell culture (Suppl. Figure 4B). Together, this data implicates ECT2 as a miR-1300 target that may play a role in mediating its effects on glioblastoma cells.

### ECT2 is a direct target of miR-1300

In order to validate *ECT2* as a direct target of miR-1300, we first transfected U251 and KNS42 cells with siRNAs directed against *ECT2* (∼95% reduction in U251 and ∼80% reduction in KNS42 (Figure 3D and Suppl. Figure 5B)). Figures 3A and 3B show that *ECT2* siRNA replicated the binucleated phenotype induced by miR-1300 at an equivalent 72 hours’ time point previously observed for mimic miR-1300. Transfection of U251 with miR-1300 caused approximately 30% and 7% increase in bi- and multi-nuclear cells respectively. Consistent with identification of ECT2 as a miR-1300 target transfection with miR-1300 induced a decrease in the expression of ECT2 of approximately 70% and 50% in U251 and KNS42 cells respectively (Figure 3C and Suppl. Figure 5A); siRNA mediated knock-down of *ECT2* caused an increase in bi- and multi-nucleated cells of approximately 40% and 15%, respectively. A similar trend was observed in KNS42 cells (Figure 3A and 3B). In addition, Western blotting experiments confirmed that transfection with miR-1300 induces a decrease in the expression of ECT2 of approximately 70% and 50% in U251 and KNS42 cells, respectively (Figure 3C and Suppl. Figure 5A). Experiments in the patient-derived GSCs produced the same result (Suppl. Figure 6). We did observe structural differences in actin and tubulin between miR-1300 expressing and *ECT2* knock-down cells by immunofluorescence (Figure 3B). This indicates that other miR1300 target genes, likely from the list of 21 targets we previously identified (Suppl. Table 5), are involved in its downstream phenotype.

**Figure 3:**
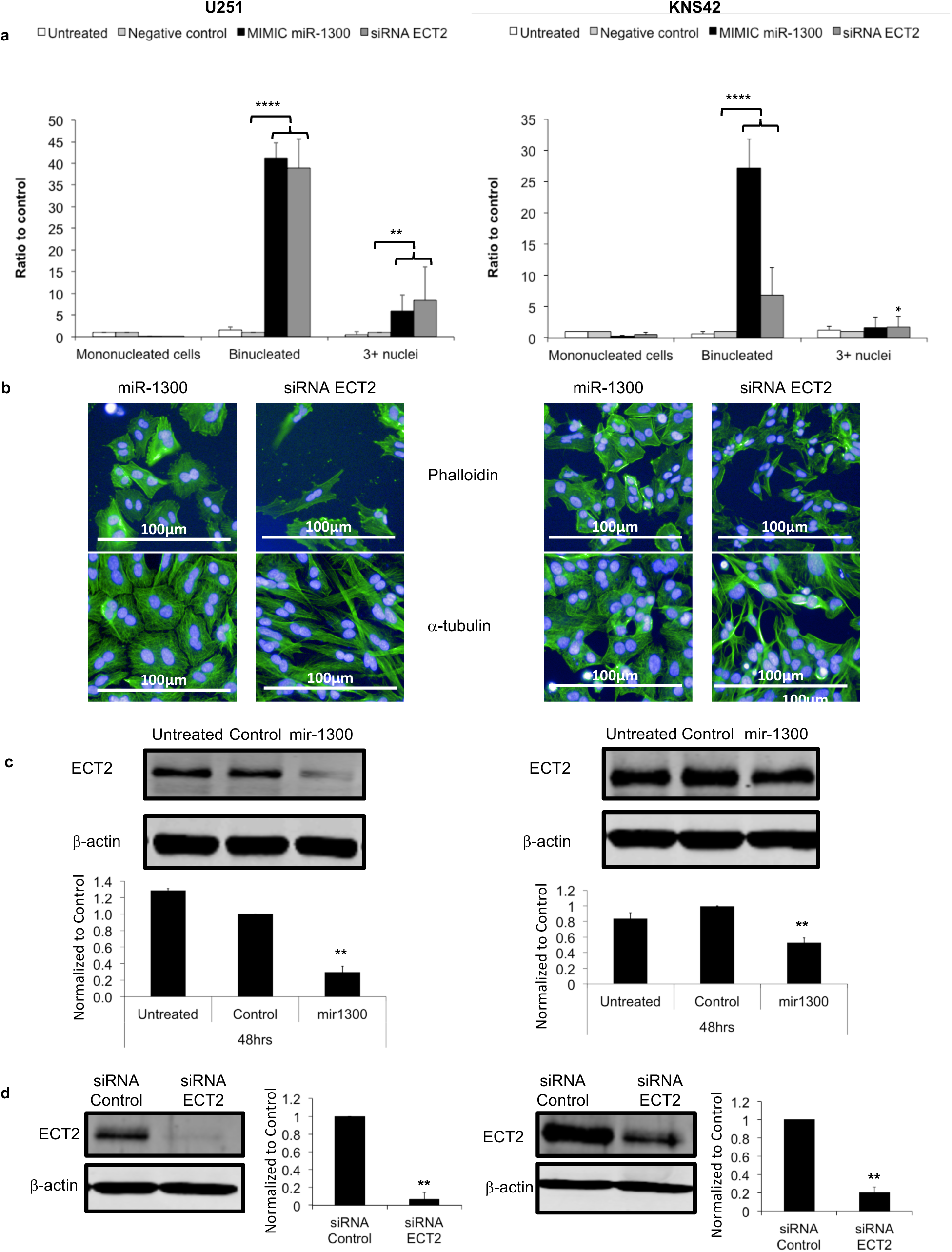
Reduced ECT2 levels in response to miR-1300 expression is associated with cytokinesis failure in U251 and KNS42 cell lines **a**: Transfection with ECT2 siRNA leads to an increase in binucleated cells. Binucleation phenotype scoring (3-6 fields of view (FOV) per condition, representing at least 100 cells) was performed on images taken on the Operetta imaging platform (x10 objective). **b**: Immunofluorescence images comparing the actin and alpha-tubulin staining in U251 and KNS42 cells 72h following transfection with 100nM of either miR-1300 or ECT2 siRNA showing both the common effect on binucleation (DAPI, blue) and the comparative differences in actin (Phalloidin) and α-tubulin structures (Alexafluor488, green). Scale bar = 100μm. **c**: Ectopic expression of miR-1300 leads to decreased expression of ECT2 at the protein level. **d**: Confirmation of ECT2 knock-down at the protein level (by WB) following transfection with ECT2 siRNA smartpool. All experiments were performed in triplicate. Results were normalized to a scrambled mimic control. Statistical significance is expressed as follows: ** = p<0.01, **** = p<0.0001. NB: Blot images were taken at 48h time point since at 72h, cells in control conditions have reached confluence, and were not expressing ECT2 anymore as they are not dividing.

Having shown that the miR-1300 phenotype is consistent amongst a range of established and patient-derived GBM lines and that *ECT2* is a promising target of miR-1300 across all cell lines tested, we went on to confirm direct targeting using 3’UTR reporter assays in the established KNS42 cell line and one patient-derived GSC culture (GBM4). Cells were transfected with a luciferase reporter containing either the wild type *ECT2* 3’UTR (3’ECT2) region or a mutated version of the *ECT2* 3’UTR (3’ECT2-mt) harboring two point mutations in the predicted miR-1300 seed region (see Methods). Co-transfection of either KNS42 or GBM4 with miR-1300 and 3’ECT2 caused a significant reduction in reporter signal in both KNS42 and GBM4 cells (Figure 4A and B, respectively). In cells transfected with 3’ECT2-mt the effect of miR-1300 on the reporter signal was abolished thus showing that the *ECT2* 3’UTR is a direct target of miR-1300 (Figure 4A and B, respectively).

**Figure 4:**
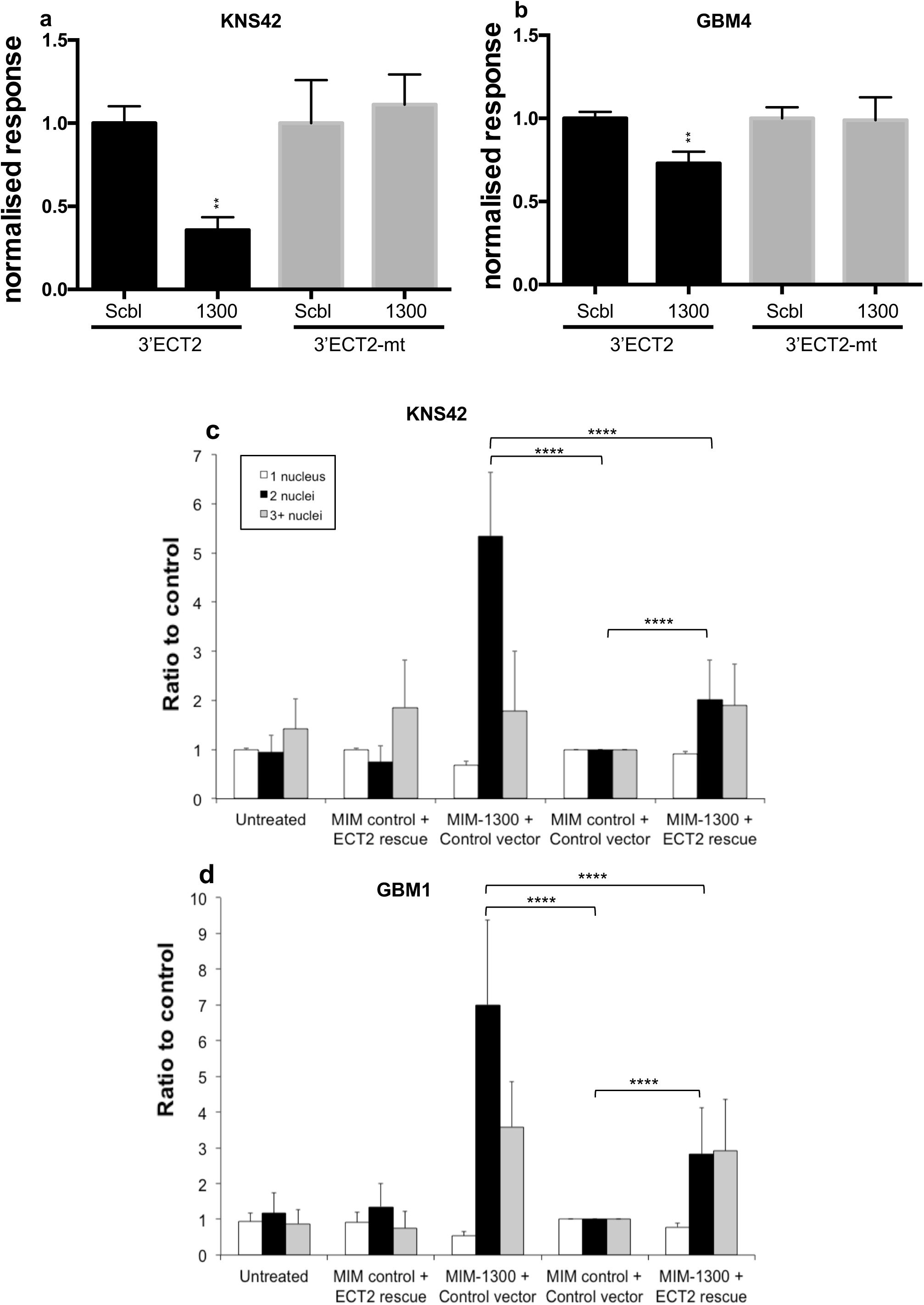
ECT2 is a direct target of miR-1300. Direct targeting assessed by 3’UTR luciferase assay in the KNS42 cells (**a**) and GBM1 GSCs (**b**). 3’ ECT2 represents the wild-type 3’UTR sequence, 3’ECT2-mt represents the 3’UTR sequence containing two point-mutations in the predicted binding site for the miR-1300 seed sequence.. C and d: Ectopic expression of ECT2 rescues miR-1300 induced cytokinesis failure. Binucleation phenotype scoring (3-6 FOV per condition, representing at least 100 cells) was performed on images taken on the Operetta imaging platform (x10 objective) in KNS42 (**c**) and GBM1 (**d**). All experiments were performed in triplicate. Results are normalized to the double control: “MIM control + Control vector” representing the scrambled mimic combined with the empty vector devoid of the ECT2 expression cassette. Statistical significance is expressed as follows: ** = p<0.01 and **** = p<0.0001.

In order to further validate the involvement of *ECT2* in the pathway downstream of miR-1300, we performed rescue experiments as follows; 36 hours following transfection with miR-1300 in KNS42 cells (∼60h in GBM1 cells), cells were transfected again with an expression vector for *ECT2* (lacking the 3’UTR region) in an attempt to rescue cells from cytokinesis failure. We showed that re-expression of ECT2 caused a 50% reduction in the number of binucleated cells in both KNS42 and GBM1 cells (Figure 4 C and D). Overall these data confirm that the effect of miR-1300 expression is mediated via reduced ECT2 levels, which drive failed cytokinesis and apoptosis in glioma cells.

### miR-1300 and ECT2 as regulator of endomitosis

Endogenous levels of *ECT2* mRNA and protein are known to be decreased during megakaryocytic differentiation at the endomitotic stage when multinucleation occurs (19). This suggests a likely regulatory role for miR-1300 on *ECT2* expression levels in platelet formation which is supported by low levels of miR-1300 remaining in platelets post-terminal differentiation (see supplementary data of ref 15) (19, 45). Using the megakaryocytic cell line CMK (21) and the Src inhibitor SU6656 which has been shown to induce endomitosis and differentiation of CMK cells into platelets (20, 21), we have now confirmed these findings and used this model of induced endomitosis to validate this previously undescribed physiological role of miR-1300. We measured a time dependent increase in expression of endogenous microRNA-1300 in CMK cells by approximately 4-fold at 48h, and approximately 6-fold at 72h and a concomitant decrease in ECT2 protein levels following exposure to 5μM SU6656 (Figure 5A and suppl. Figure 7). Moreover, we confirmed the increase of polyploid megakaryocytic cells by nearly 2-fold at 24h, and 1.5-fold both at 48h and 72h respectively, using high-content immunofluorescence imaging (Figure 5B).

### Differentiated brain tumor cells are not affected by miR-1300 expression

In order to establish whether the phenotype caused by miR-1300 was specific to glioma cells, we assessed the effect of its ectopic expression on fully differentiated GSCs, compared to their stem-like, proliferative (isogenic) counterparts. Differentiation of GBM1 and GBM4 cells was achieved by 5 days exposure to BMP4 as previously described (22-24, 46, 47).

In GBM1 cells, transfection with miR-1300 caused a 4-fold increase in the proportion of cells in G2/M phase, which was significantly reduced by 2.3-fold in the BMP4 differentiated counterparts (Figure 6A). In addition, miR-1300 caused a 1.5-fold reduction in live cell numbers, a 3.8-fold increase in early apoptotic cells and a 14-fold increase in apoptotic cells, whereas there was no significant change in their differentiated counterparts (Figure 6B). We also observed a reduction by half in the number of binucleated cells in BMP4 differentiated GSCs compared to non-differentiated cells. GBM4 cells showed a 0.8-fold increase in cells in G2/M after transfection with miR-1300, which was reduced to a 0.2-fold after BMP4 treatment (Figure 6A). MiR-1300 caused a 50% reduction in live GBM4 cells and a 3.8 and 4-fold increase in early-mid apoptotic and mid-late apoptotic cells, respectively. However, cell death was significantly reduced in BMP4 treated post mitotic GBM4 cells transfected with miR-1300, to the point where there were no significant differences in comparison with the scrambled control transfected cells (Figure 6B). In addition, BMP4 treated GBM4 cells did not show an increased number of binucleated cells compared to control cells, as opposed to their non-differentiated counterparts (Figure 7B). Taken together these data suggest that expression of miR-1300 in non-proliferating, differentiated cells does not induce significant cell cycle arrest or apoptosis when compared to proliferating, undifferentiated (stem-like) isogenic pairs. This suggests that miR-1300 could represent an attractive target with a favorable therapeutic ratio in glioma.

**Figure 6:**
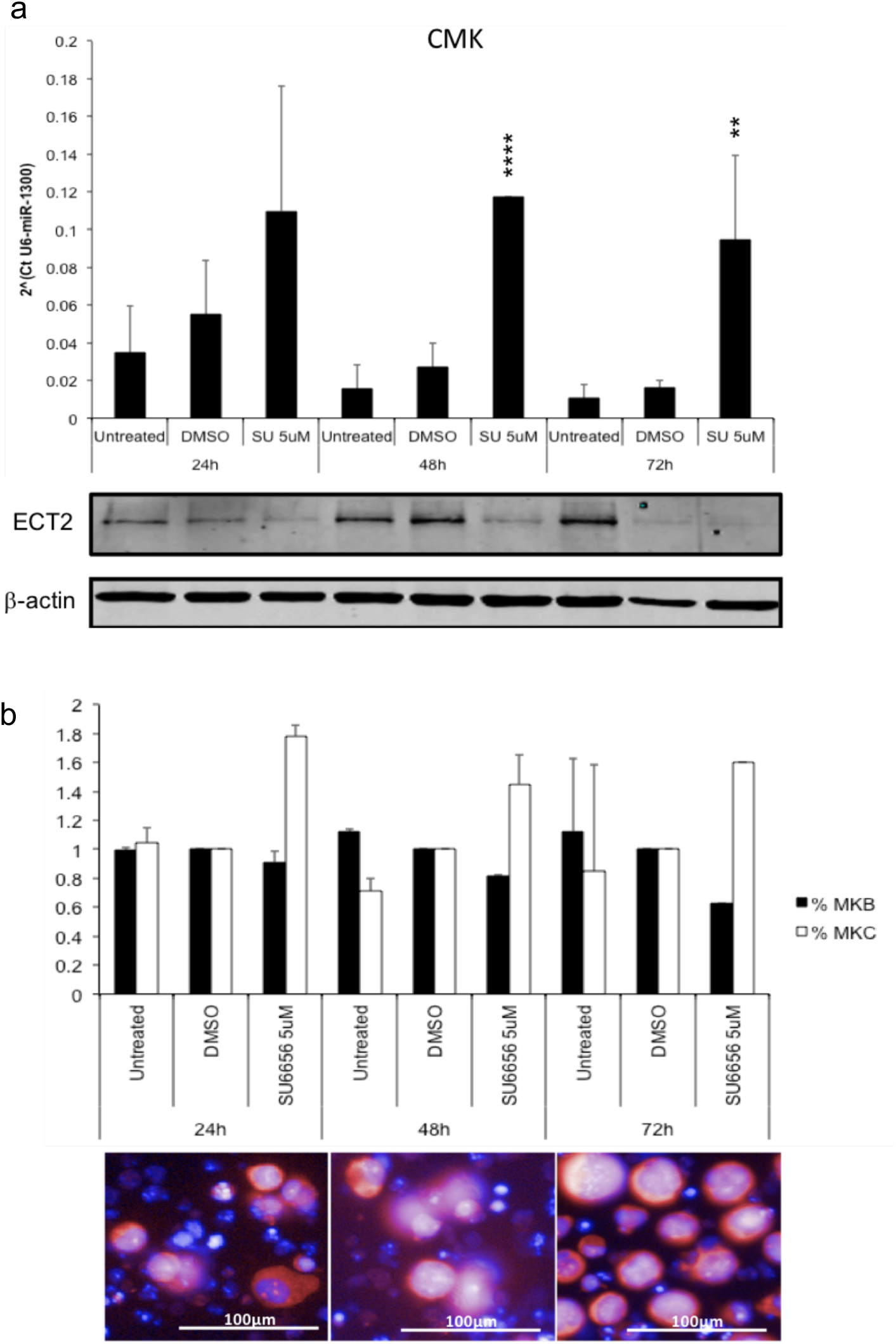
Ectopic expression of miR-1300 specifically affects stem-like cells but not their differentiated counterparts. (**a**) Effect on the cell cycle was analyzed by flow cytometry 120h post-transfection using Propidium Iodide (PI) loading as a measure of DNA content. (**b**) Cell death measured also 120h post-transfection by flow cytometry where Annexin negative (An -), PI negative (PI-) = live cells, An+ PI- = early apoptotic cells, An+ PI+ = mid/late apoptotic cells and An-PI+ = Necrotic cells. (**c**) Binucleation phenotype scoring (3-6 images per condition, representing at least 100 cells) was performed on images taken on the Operetta imaging platform (x10). All experiments were performed in triplicate. Results are normalized to a scrambled mimic control without BMP4 exposure. Statistical significance is expressed as follows: * = p<0.05, ** = p<0.01, *** = p<0.001 and **** = p<0.0001.

**Figure 7:**
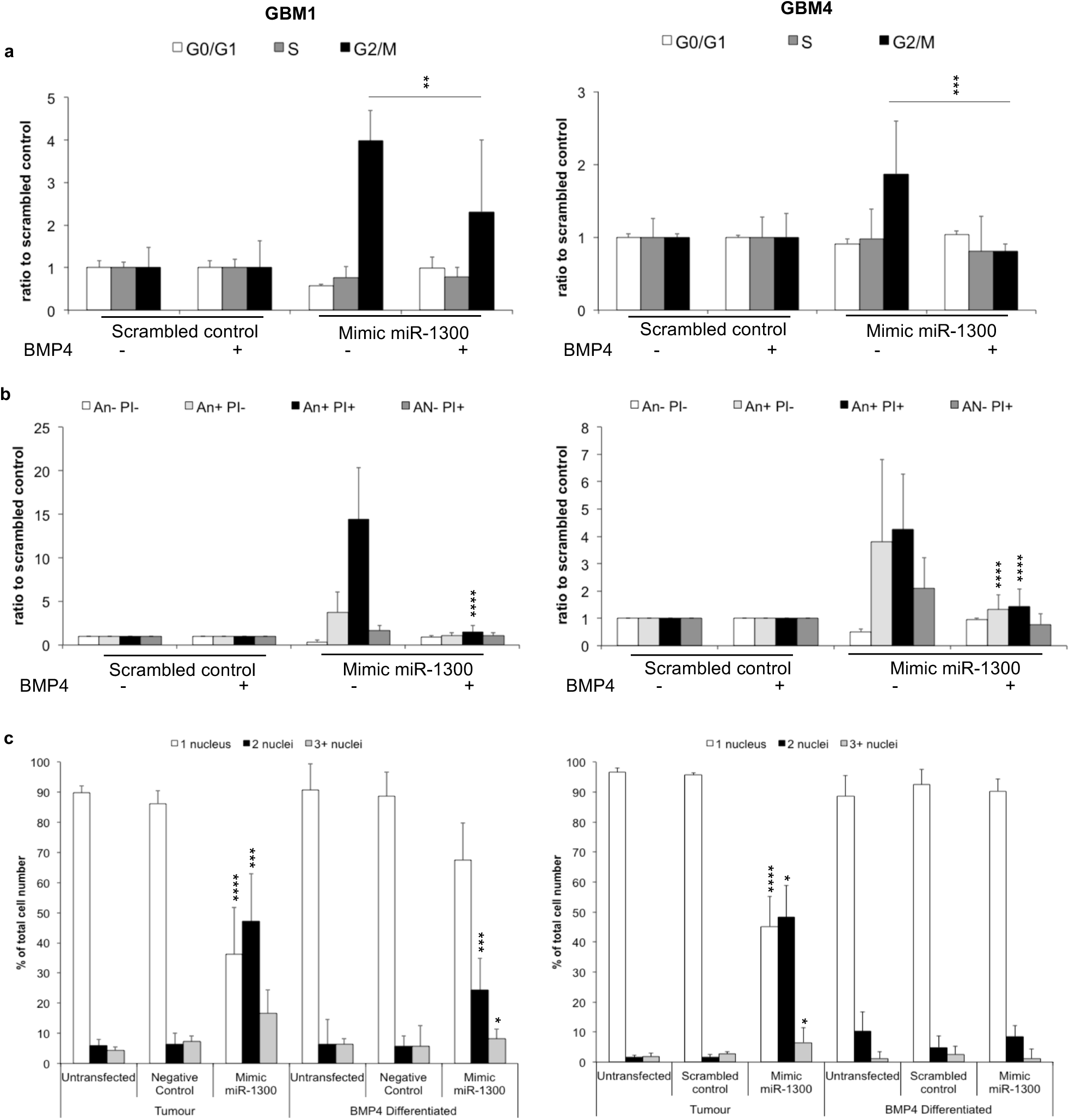
Ectopic expression of miR-1300 specifically affects stem-like cells but not their differentiated counterparts

## Conclusions

Our high-throughput screen defined the landscape of microRNAs with significant cytotoxic effects in adult and pediatric GBM cell lines.

We have shown that ectopic expression of miR-1300, the most potent candidate highlighted by the screen, leads to cytokinesis failure followed by apoptosis in both established and in patient-derived GBM cells, importantly, without affecting terminally differentiated glioma cells. Finally, we have validated ECT2 as one of the direct effectors downstream of miR-1300.

The effect of miR-1300 shows its translational potential as a treatment for GBM. It also makes it a very promising candidate for combination therapy as a chemo-radiosensitizer decreasing the ability of dividing cells to recover from the damage induced by conventional therapy. Moreover, this effect could be of paramount use in the second line of treatment where by specifically targeting resistant repopulating cells it could also significantly impair recurrence.

Ongoing work is aimed at validating the mechanism of action of miR-1300 in more detail as well as addressing delivery of microRNAs for brain tumor treatment and its potential use in combination with conventional treatments.

## Acknowledgements

Dr Adam Davison and Miss Liz Straszynski from the Flow-Cytometry and Imaging Facility provided training, advice and analysis support for all confocal microscopy and flow-cytometry based assays. Dr Claire Taylor authenticated the cell lines. Miss Christina Ndidi Efua Okafor performed a review of the literature. Dr Georgia Mavria provided support and guidance in the writing of the manuscript and interpretation of the cytokinesis data.

## List of abbreviations

GBM: Glioblastoma
GSC: Glioblastoma Stem Cell
An: AnnexinV-FITC
PI: Propidium Iodide
Scbl: Scramble (non-targeting) microRNA control
GEF: GTP Exchange Factor
*ECT2*: Epithelial Cell Transforming 2
miR-1300: mimic microRNA-1300
ORF: Open Reading Frame
GO: Gene Ontology
MKB: megakaryoblast
MKC: Megakaryocyte

## Figure Captions

Exposure of synchronised CMK cells to SU6656 concomitantly induces an increase in the levels of miR-1300 and a decrease of its targets, ECT2 (**a**). Cells were synchronised using Monastrol 25μM for 24h prior to exposure to SU6656 5μM. (**b**) Increase in multinuclear megakaryocytic cells (MKC) as a result of endomitosis undergone by megakaryoblastic cells (MKB) was measured by immunofluorescence in response to SU6656. Cells were stained with DAPI and TOTO-3. Nine fields of view were analysed using an algorithm designed in the Columbus software to discriminate objects based on their size (**b**). Statistical significance is expressed as follows: * = p<0.05, ** = p<0.01, *** = p<0.001 and **** = p<0.0001.

