## Supplementary files for "A high-content screen profiles cytotoxic microRNAs in pediatric and adult glioblastoma cells and identifies miR-1300 as a potent inducer of cytokinesis failure"

**Additional files (in order of appearance in the text)**

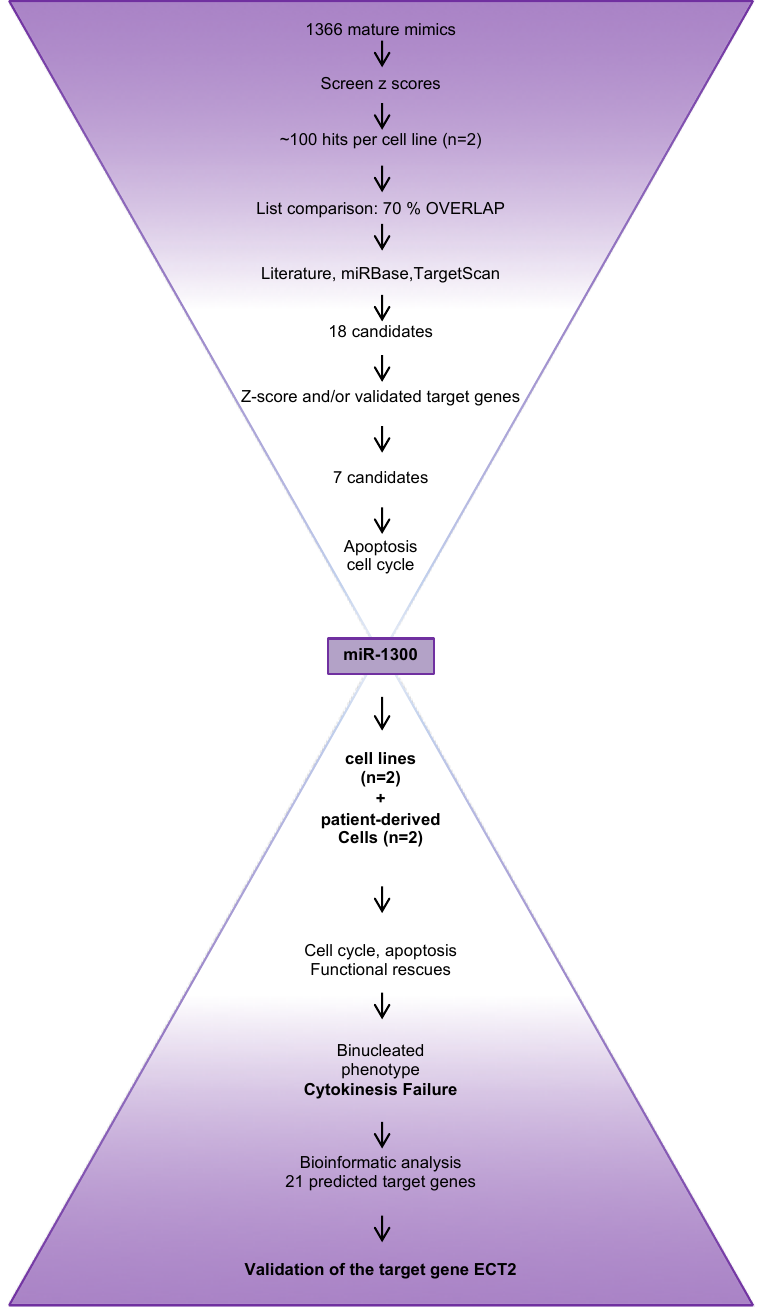

Supplementary Figure1: Project overview: Key steps of the project from the high-throughput screen to the final target validation

| **U251** | |  | **KNS42** | |  |
| --- | --- | --- | --- | --- | --- |
| **zb mean** | **microRNA** |  | **zbmean** | **microRNA** | **also a hit in U251** |
| **-5.14** | **hsa-miR-1300** |  | -5.31 | hsa-miR-30b*= 30b-3p | ✓ |
| -5.01 | hsa-miR-876-3p |  | -5.23 | hsa-miR-1294 | ✓ |
| -4.99 | hsa-miR-516b=516b-5p |  | -4.74 | hsa-miR-644=644a | ✓ |
| -4.95 | hsa-miR-3115 |  | -4.53 | hsa-miR-541=541-3p | ✓ |
| -4.84 | hsa-miR-3165 |  | -4.39 | hsa-miR-380*=380-5p |  |
| -4.79 | hsa-miR-3140=3140-3p |  | -4.34 | hsa-miR-1293 | ✓ |
| -4.61 | hsa-miR-1293 |  | -4.29 | hsa-miR-765 | ✓ |
| -4.31 | hsa-miR-765 |  | -4.20 | hsa-miR-885-3p | ✓ |
| -4.24 | hsa-miR-3691=3691-5p |  | -4.18 | hsa-let-7g*= let-7g-3p | ✓ |
| -4.19 | hsa-miR-205*=205-3p |  | -4.14 | hsa-miR-449a | ✓ |
| -4.16 | hsa-miR-216a=216a-5p |  | -3.99 | hsa-miR-1289 | ✓ |
| -4.15 | hsa-miR-1244 |  | -3.96 | hsa-miR-3907 | ✓ |
| -4.10 | hsa-miR-184 |  | -3.95 | hsa-miR-3689b*=3689-3p | ✓ |
| -4.09 | hsa-miR-613 |  | -3.94 | hsa-miR-1269 | ✓ |
| -4.08 | hsa-miR-3158=3158-3p |  | -3.89 | hsa-miR-34a=34a-5p | ✓ |
| -4.05 | hsa-miR-671-5p |  | -3.82 | hsa-miR-876-3p | ✓ |
| -4.04 | hsa-miR-185*=185-3p |  | -3.81 | hsa-miR-3936 | ✓ |
| -4.04 | hsa-miR-3689b*=3689b-3p |  | -3.76 | hsa-miR-1306 | ✓ |
| -4.02 | hsa-miR-369-5p |  | -3.67 | hsa-miR-558 | ✓ |
| -3.99 | hsa-miR-3916 |  | -3.66 | hsa-miR-432*=432-3p | ✓ |
| -3.98 | hsa-miR-147=147a |  | -3.65 | hsa-miR-1197 | ✓ |
| -3.97 | hsa-miR-486-3p |  | -3.55 | hsa-miR-593=mir-593-3p | ✓ |
| -3.93 | hsa-miR-3154 |  | -3.53 | hsa-miR-1275 | ✓ |
| -3.92 | hsa-miR-4306 |  | -3.46 | hsa-miR-453=323b-5p |  |
| -3.91 | hsa-miR-449b=449b-5p |  | -3.45 | hsa-miR-891b | ✓ |
| -3.90 | hsa-miR-449c= 449c-5p |  | -3.44 | hsa-miR-1827 | ✓ |
| -3.83 | hsa-miR-665 |  | -3.40 | hsa-miR-604 | ✓ |
| -3.81 | hsa-miR-149*=149-3p |  | -3.35 | hsa-miR-665 | ✓ |
| -3.76 | hsa-miR-3179 |  | -3.29 | hsa-miR-34c-5p | ✓ |
| -3.74 | hsa-miR-3160=3160-3p |  | -3.26 | hsa-miR-609 |  |
| -3.74 | hsa-miR-4297 |  | -3.26 | hsa-miR-3140-3p | ✓ |
| -3.72 | hsa-miR-647 |  | -3.25 | hsa-miR-549 |  |
| -3.72 | hsa-miR-555 |  | -3.23 | hsa-miR-193b*=193-5p |  |
| -3.65 | hsa-miR-520d-5p=520d* |  | -3.16 | hsa-miR-3612 |  |
| -3.65 | hsa-miR-127-5p |  | -3.15 | hsa-miR-802 | ✓ |
| -3.64 | hsa-miR-3918 |  | -3.15 | hsa-miR-3160= 3160-3p | ✓ |
| -3.63 | hsa-miR-940 |  | -3.14 | hsa-miR-621 |  |
| -3.61 | hsa-miR-34c-5p |  | -3.07 | hsa-miR-637 | ✓ |
| -3.60 | hsa-miR-1321 |  | -3.06 | hsa-miR-671-3p | ✓ |
| -3.59 | hsa-miR-34b*=5p |  | -3.06 | hsa-miR-365=hsa-miR-365a-3p =hsa-miR-365b-3p |  |
| -3.55 | hsa-miR-3168 |  | -3.01 | hsa-miR-3124=3124-5p | ✓ |
| -3.52 | hsa-miR-604 |  | -3.00 | hsa-miR-3911 | ✓ |
| -3.51 | hsa-miR-142-3p |  | **-2.98** | **hsa-miR-1300** | ✓ |
| -3.51 | hsa-miR-885-3p |  | -2.97 | hsa-miR-555 | ✓ |
| -3.50 | hsa-miR-449a |  | -2.96 | hsa-miR-1250 |  |
| -3.47 | hsa-miR-624=624-3p |  | -2.87 | hsa-miR-3192 | ✓ |
| -3.45 | hsa-miR-3675-5p |  | -2.82 | hsa-miR-2110 | ✓ |
| -3.40 | hsa-miR-617 |  | -2.79 | hsa-miR-647 | ✓ |
| -3.37 | hsa-miR-1909=1909-3p |  | -2.76 | hsa-miR-342-5p | ✓ |
| -3.36 | hsa-miR-1244 |  | -2.75 | hsa-miR-3186-3p | ✓ |
| -3.35 | hsa-miR-148b*=148b-5p |  | -2.74 | hsa-miR-1205 | ✓ |
| -3.34 | hsa-miR-124=124-3p |  | -2.73 | hsa-miR-624 | ✓ |
| -3.34 | hsa-miR-509-3-5p |  | -2.72 | hsa-miR-892b |  |
| -3.33 | hsa-miR-3677=3677-3p |  | -2.72 | hsa-miR-491-5p | ✓ |
| -3.31 | hsa-miR-340=340-5p |  | -2.69 | hsa-miR-299-3p |  |
| -3.28 | hsa-miR-509-3p |  | -2.67 | hsa-miR-329 |  |
| -3.27 | hsa-miR-2909 |  | -2.67 | hsa-miR-520d-5p | ✓ |
| -3.26 | hsa-miR-1263 |  | -2.66 | hsa-miR-650 | ✓ |
| -3.23 | hsa-let-7g*= let-7g-3p |  | -2.64 | hsa-miR-4318 |  |
| -3.23 | hsa-miR-518d-3p |  | -2.63 | hsa-miR-22*=22-5p |  |
| -3.21 | hsa-miR-3157=3175-5p |  | -2.62 | hsa-miR-193b | ✓ |
| -3.21 | hsa-miR-323-5p=323a-5p |  | -2.57 | hsa-miR-516b=5p | ✓ |
| -3.21 | hsa-miR-449c*=449c-3p |  | -2.56 | hsa-miR-506 | ✓ |
| -3.19 | hsa-miR-200b=200b-3p |  | -2.55 | hsa-miR-323-5p=323a-5p | ✓ |
| -3.19 | hsa-miR-363*=363-5p |  | -2.54 | hsa-miR-340=340-5p | ✓ |
| -3.19 | hsa-miR-548aa |  | -2.54 | hsa-miR-34b*=34b-5p | ✓ |
| -3.18 | hsa-miR-1270 |  | -2.53 | hsa-miR-296-5p |  |
| -3.18 | hsa-miR-3169 |  | -2.46 | hsa-miR-4316 |  |
| -3.16 | hsa-miR-25*=25-5p |  | -2.45 | hsa-miR-585 |  |
| -3.16 | hsa-miR-744=744-5p |  | -2.44 | hsa-miR-542-3p |  |
| -3.15 | hsa-miR-634 |  | -2.43 | hsa-miR-4283 |  |
| -3.15 | hsa-miR-650 |  | -2.41 | hsa-let-7a-2*=hsa-let-7a-2-3p | ✓ |
| -3.14 | hsa-miR-208a |  | -2.40 | hsa-miR-214*=hsa-miR-214-5p | ✓ |
| -3.14 | hsa-miR-493*=493-5p |  | -2.40 | hsa-miR-10b=10b-5p |  |
| -3.13 | hsa-miR-101*=101-5p |  | -2.40 | hsa-miR-449b=449b-5p | ✓ |
| -3.12 | hsa-miR-302b=302b-3p |  | -2.36 | hsa-miR-3119 | ✓ |
| -3.12 | hsa-miR-3173=3173-3p |  | -2.34 | hsa-miR-3179 | ✓ |
| -3.12 | hsa-miR-3186-3p |  | 2.39 | hsa-miR-1277 |  |
| -3.10 | hsa-miR-34a=34a-5p |  | 2.58 | hsa-miR-1280 |  |
| -3.10 | hsa-miR-185=185-5p |  | 2.90 | hsa-miR-625*=625-3p |  |
| -3.09 | hsa-miR-891b |  | 3.21 | hsa-miR-199a-5p | ✓ |
| -3.09 | hsa-miR-597 |  | 3.28 | hsa-miR-375 |  |
| -3.09 | hsa-miR-644=644a |  | 3.96 | hsa-miR-23a |  |
| -3.09 | hsa-miR-671-3p |  |  |  |  |
| -3.08 | hsa-miR-502-3p |  |  |  |  |
| -3.08 | hsa-miR-1193 |  |  |  |  |
| -3.08 | hsa-miR-3116 |  |  |  |  |
| -3.06 | hsa-miR-3126-5p |  |  |  |  |
| -3.06 | hsa-miR-3189=3189-3p |  |  |  |  |
| -3.06 | hsa-miR-1322 |  |  |  |  |
| -3.05 | hsa-miR-3662 |  |  |  |  |
| -3.05 | hsa-miR-3647-5p |  |  |  |  |
| -3.05 | hsa-miR-567 |  |  |  |  |
| -3.05 | hsa-miR-302a=302a-3p |  |  |  |  |
| -3.05 | hsa-miR-3653 |  |  |  |  |
| -3.04 | hsa-miR-3177=3177-3p |  |  |  |  |
| -3.04 | hsa-miR-491-5p |  |  |  |  |
| -3.03 | hsa-miR-3619=3619-5p |  |  |  |  |
| -3.02 | hsa-miR-520a-5p=520a* |  |  |  |  |
| -3.02 | hsa-miR-3119 |  |  |  |  |
| -3.01 | hsa-let-7f-1*=let-7f-1-3p |  |  |  |  |
| -3.01 | hsa-miR-103-as=103b |  |  |  |  |
| -3.01 | hsa-miR-3923 |  |  |  |  |
| -3.00 | hsa-miR-637 |  |  |  |  |
| -3.00 | hsa-miR-3622b-5p |  |  |  |  |
| -3.00 | hsa-miR-3200-3p |  |  |  |  |
| -2.99 | hsa-miR-330-5p |  |  |  |  |
| -2.98 | hsa-miR-432* |  |  |  |  |
| -2.97 | hsa-miR-378* |  |  |  |  |
| -2.96 | hsa-miR-101=101-3p |  |  |  |  |
| -2.95 | hsa-miR-1200 |  |  |  |  |
| -2.95 | hsa-miR-138 |  |  |  |  |
| -2.94 | hsa-miR-654-5p |  |  |  |  |
| -2.92 | hsa-miR-129-5p |  |  |  |  |
| -2.92 | hsa-miR-16-1* |  |  |  |  |
| -2.91 | hsa-miR-1275 |  |  |  |  |
| -2.90 | hsa-miR-30b* |  |  |  |  |
| -2.90 | hsa-miR-18b |  |  |  |  |
| -2.90 | hsa-miR-219-1-3p |  |  |  |  |
| -2.87 | hsa-miR-605 |  |  |  |  |
| -2.87 | hsa-miR-556-3p |  |  |  |  |
| -2.87 | hsa-miR-500* |  |  |  |  |
| -2.86 | hsa-miR-383 |  |  |  |  |
| -2.84 | hsa-miR-508-3p |  |  |  |  |
| -2.83 | hsa-miR-3924 |  |  |  |  |
| -2.83 | hsa-miR-3191 |  |  |  |  |
| -2.83 | hsa-miR-2115* |  |  |  |  |
| -2.82 | hsa-miR-376b |  |  |  |  |
| -2.81 | hsa-miR-1272 |  |  |  |  |
| -2.81 | hsa-miR-601 |  |  |  |  |
| -2.81 | hsa-miR-3907 |  |  |  |  |
| -2.80 | hsa-miR-3161 |  |  |  |  |
| -2.79 | hsa-miR-193b |  |  |  |  |
| -2.79 | hsa-miR-2278 |  |  |  |  |
| -2.78 | hsa-miR-324-5p |  |  |  |  |
| -2.78 | hsa-miR-3616-3p |  |  |  |  |
| -2.77 | hsa-miR-3910 |  |  |  |  |
| -2.77 | hsa-miR-3673 |  |  |  |  |
| -2.75 | hsa-miR-576-3p |  |  |  |  |
| -2.75 | hsa-miR-1 |  |  |  |  |
| -2.75 | hsa-miR-1304 |  |  |  |  |
| -2.73 | hsa-miR-648 |  |  |  |  |
| -2.73 | hsa-miR-608 |  |  |  |  |
| -2.73 | hsa-miR-181c* |  |  |  |  |
| -2.72 | hsa-miR-196a* |  |  |  |  |
| -2.71 | hsa-miR-1270 |  |  |  |  |
| -2.71 | hsa-miR-1288 |  |  |  |  |
| -2.70 | hsa-miR-370 |  |  |  |  |
| -2.70 | hsa-let-7a-2* |  |  |  |  |
| -2.70 | hsa-miR-1294 |  |  |  |  |
| -2.70 | hsa-miR-2110 |  |  |  |  |
| -2.69 | hsa-miR-151-3p |  |  |  |  |
| -2.69 | hsa-miR-3117 |  |  |  |  |
| -2.69 | hsa-miR-378 |  |  |  |  |
| -2.68 | hsa-miR-3170 |  |  |  |  |
| -2.68 | hsa-miR-192* |  |  |  |  |
| -2.67 | hsa-miR-219-2-3p |  |  |  |  |
| -2.67 | hsa-miR-3663-3p |  |  |  |  |
| -2.66 | hsa-miR-138-2* |  |  |  |  |
| -2.64 | hsa-miR-924 |  |  |  |  |
| -2.62 | hsa-miR-3650 |  |  |  |  |
| -2.62 | hsa-miR-1827 |  |  |  |  |
| -2.61 | hsa-miR-1269 |  |  |  |  |
| -2.61 | hsa-miR-1273e |  |  |  |  |
| -2.60 | hsa-miR-520c-3p |  |  |  |  |
| -2.60 | hsa-miR-3121 |  |  |  |  |
| -2.59 | hsa-miR-1295 |  |  |  |  |
| -2.59 | hsa-miR-544 |  |  |  |  |
| -2.59 | hsa-miR-3118 |  |  |  |  |
| -2.59 | hsa-miR-1289 |  |  |  |  |
| -2.59 | hsa-miR-518a-5p |  |  |  |  |
| -2.58 | hsa-miR-3672 |  |  |  |  |
| -2.58 | hsa-miR-3132 |  |  |  |  |
| -2.57 | hsa-miR-3180-3p |  |  |  |  |
| -2.56 | hsa-miR-206 |  |  |  |  |
| -2.56 | hsa-miR-541 |  |  |  |  |
| -2.56 | hsa-miR-374c |  |  |  |  |
| -2.56 | hsa-miR-3936 |  |  |  |  |
| -2.55 | hsa-miR-28-3p |  |  |  |  |
| -2.54 | hsa-miR-630 |  |  |  |  |
| -2.54 | hsa-miR-135b* |  |  |  |  |
| -2.53 | hsa-miR-625 |  |  |  |  |
| -2.52 | hsa-miR-3616-5p |  |  |  |  |
| -2.52 | hsa-miR-376a* |  |  |  |  |
| -2.52 | hsa-miR-22 |  |  |  |  |
| -2.50 | hsa-miR-3135 |  |  |  |  |
| -2.50 | hsa-miR-3689a-3p |  |  |  |  |
| -2.50 | hsa-miR-506 |  |  |  |  |
| -2.49 | hsa-miR-3127 |  |  |  |  |
| -2.48 | hsa-miR-520b |  |  |  |  |
| -2.48 | hsa-miR-3192 |  |  |  |  |
| -2.48 | hsa-miR-9* |  |  |  |  |
| -2.47 | hsa-miR-105 |  |  |  |  |
| -2.46 | hsa-miR-208b |  |  |  |  |
| -2.46 | hsa-miR-548s |  |  |  |  |
| -2.44 | hsa-miR-202 |  |  |  |  |
| -2.43 | hsa-miR-802 |  |  |  |  |
| -2.42 | hsa-miR-19b-1* |  |  |  |  |
| -2.42 | hsa-miR-548q |  |  |  |  |
| -2.42 | hsa-miR-1972 |  |  |  |  |
| -2.42 | hsa-miR-3139 |  |  |  |  |
| -2.41 | hsa-miR-126 |  |  |  |  |
| -2.40 | hsa-miR-145* |  |  |  |  |
| -2.40 | hsa-miR-3682 |  |  |  |  |
| -2.39 | hsa-miR-561 |  |  |  |  |
| -2.38 | hsa-miR-1282 |  |  |  |  |
| -2.37 | hsa-miR-657 |  |  |  |  |
| -2.37 | hsa-miR-3675-3p |  |  |  |  |
| -2.37 | hsa-miR-92a-1* |  |  |  |  |
| -2.37 | hsa-miR-371-3p |  |  |  |  |
| -2.36 | hsa-miR-624* |  |  |  |  |
| -2.36 | hsa-miR-20b* |  |  |  |  |
| -2.36 | hsa-miR-1205 |  |  |  |  |
| -2.35 | hsa-miR-2114* |  |  |  |  |
| -2.35 | hsa-miR-764 |  |  |  |  |
| -2.34 | hsa-miR-3146 |  |  |  |  |
| -2.34 | hsa-miR-3142 |  |  |  |  |
| -2.34 | hsa-miR-766 |  |  |  |  |
| -2.33 | hsa-miR-342-5p |  |  |  |  |
| -2.31 | hsa-miR-3714 |  |  |  |  |
| -2.31 | hsa-miR-3652 |  |  |  |  |
| -2.30 | hsa-miR-614 |  |  |  |  |
| -2.30 | hsa-miR-558 |  |  |  |  |
| -2.29 | hsa-miR-1306 |  |  |  |  |
| -2.28 | hsa-miR-593=593-3p |  |  |  |  |
| -2.28 | hsa-miR-214* |  |  |  |  |
| -2.27 | hsa-miR-499-3p |  |  |  |  |
| -2.27 | hsa-miR-525-5p |  |  |  |  |
| -2.27 | hsa-miR-3174 |  |  |  |  |
| -2.27 | hsa-miR-29a |  |  |  |  |
| -2.26 | hsa-miR-320d |  |  |  |  |
| -2.26 | hsa-miR-3137 |  |  |  |  |
| -2.26 | hsa-miR-1271 |  |  |  |  |
| -2.24 | hsa-miR-200a |  |  |  |  |
| -2.24 | hsa-miR-544b |  |  |  |  |
| -2.24 | hsa-miR-1207-5p |  |  |  |  |
| -2.24 | hsa-miR-3202 |  |  |  |  |
| -2.24 | hsa-miR-3129 |  |  |  |  |
| -2.23 | hsa-miR-452* |  |  |  |  |
| -2.23 | hsa-miR-612 |  |  |  |  |
| -2.23 | hsa-miR-512-5p |  |  |  |  |
| -2.23 | hsa-miR-382 |  |  |  |  |
| -2.22 | hsa-miR-3622a-5p |  |  |  |  |
| -2.22 | hsa-miR-129* |  |  |  |  |
| -2.21 | hsa-miR-376a |  |  |  |  |
| -2.21 | hsa-miR-3145 |  |  |  |  |
| -2.21 | hsa-miR-3186-5p |  |  |  |  |
| -2.21 | hsa-miR-146a |  |  |  |  |
| -2.20 | hsa-miR-3124=3124-5p |  |  |  |  |
| -2.20 | hsa-miR-1245 |  |  |  |  |
| -2.20 | hsa-miR-18a |  |  |  |  |
| -2.17 | hsa-miR-1178 |  |  |  |  |
| -2.16 | hsa-miR-520f |  |  |  |  |
| -2.15 | hsa-miR-1197 |  |  |  |  |
| -2.15 | hsa-miR-2117 |  |  |  |  |
| -2.14 | hsa-miR-1302 |  |  |  |  |
| -2.12 | hsa-miR-3915 |  |  |  |  |
| -2.12 | hsa-miR-488 |  |  |  |  |
| -2.11 | hsa-miR-3911 |  |  |  |  |
| -2.11 | hsa-miR-562 |  |  |  |  |
| -2.09 | hsa-miR-557 |  |  |  |  |
| -2.08 | hsa-miR-3680* |  |  |  |  |
| -2.07 | hsa-miR-30a* |  |  |  |  |
| -2.06 | hsa-miR-297 |  |  |  |  |
| -2.06 | hsa-miR-133a |  |  |  |  |
| -2.06 | hsa-miR-3649 |  |  |  |  |
| -2.06 | hsa-miR-1262 |  |  |  |  |
| -2.06 | hsa-miR-1972 |  |  |  |  |
| -2.05 | hsa-miR-3909 |  |  |  |  |
| -2.05 | hsa-miR-507 |  |  |  |  |
| -2.04 | hsa-miR-628-5p |  |  |  |  |
| -2.04 | hsa-miR-34a* |  |  |  |  |
| -2.04 | hsa-miR-573 |  |  |  |  |
| -2.04 | hsa-miR-193a-3p |  |  |  |  |
| -2.03 | hsa-miR-1296 |  |  |  |  |
| -2.03 | hsa-miR-182 |  |  |  |  |
| -2.03 | hsa-miR-3679-3p |  |  |  |  |
| -2.03 | hsa-miR-1256 |  |  |  |  |
| -2.02 | hsa-miR-1185 |  |  |  |  |
| -2.02 | hsa-miR-1298 |  |  |  |  |
| -2.01 | hsa-miR-144 |  |  |  |  |
| -2.01 | hsa-miR-302d |  |  |  |  |
| -2.00 | hsa-miR-27a* |  |  |  |  |
| 2.04 | hsa-miR-587 |  |  |  |  |
| 2.06 | hsa-miR-200c* |  |  |  |  |
| 2.06 | hsa-miR-1974 |  |  |  |  |
| 2.08 | hsa-miR-933 |  |  |  |  |
| 2.13 | hsa-miR-3684 |  |  |  |  |
| 2.14 | hsa-miR-302d* |  |  |  |  |
| 2.14 | hsa-miR-99a* |  |  |  |  |
| 2.15 | hsa-miR-602 |  |  |  |  |
| 2.16 | hsa-miR-643 |  |  |  |  |
| 2.24 | hsa-miR-300 |  |  |  |  |
| 2.24 | hsa-miR-1297 |  |  |  |  |
| 2.29 | hsa-miR-26b |  |  |  |  |
| 2.36 | hsa-miR-3664 |  |  |  |  |
| 2.39 | hsa-miR-663 |  |  |  |  |
| 2.53 | hsa-miR-187 |  |  |  |  |
| 2.54 | hsa-miR-29c* |  |  |  |  |
| 2.55 | hsa-miR-9 |  |  |  |  |
| 2.94 | hsa-miR-545=545-3p |  |  |  |  |
| 3.53 | hsa-miR-1305 |  |  |  |  |
| 3.71 | hsa-miR-1973 |  |  |  |  |
| 3.94 | hsa-miR-199a-5p |  |  |  |  |

Supplementary Table 1 Summary of all microRNA mimics causing a significant decrease, zb is the mean z score of the duplicate screens (> -2SD for KNS42 and -3SD for U251, respectively) or increase (< +2SD for KNS42 and < +3SD for U251, respectively). A tick mark “√” indicates microRNA hits present in both the adult cell line U251 and the Pediatric cell line KNS42. The hit miR-1300 is highlighted in grey in both the column of KNS42 and U251 results.

Supplementary Table 2 (KNS42 duplicate library screen results) and Table 3 (U251 duplicate library screen results) are large data tables and thus have to be seen separately.

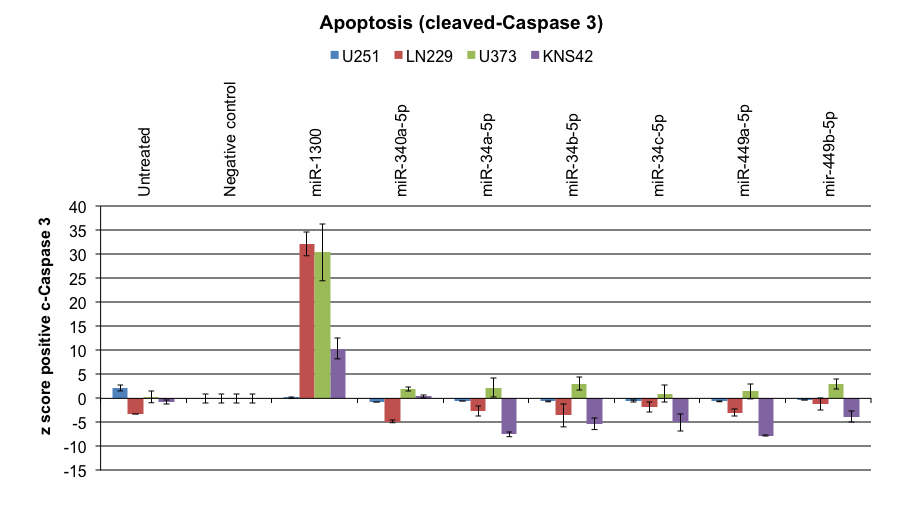

**b***:* Apoptosis

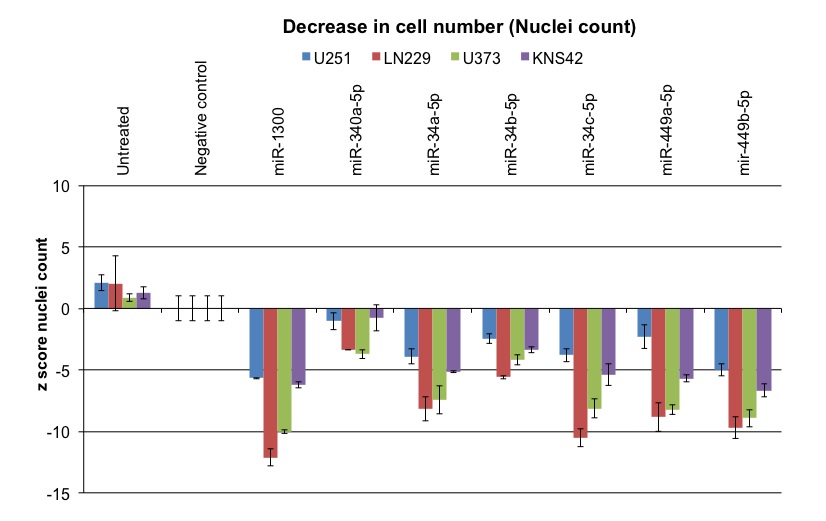

**c**: Cleaved-Caspase-3 foci, example of U251 and KNS42

**a**: Cytotoxicity

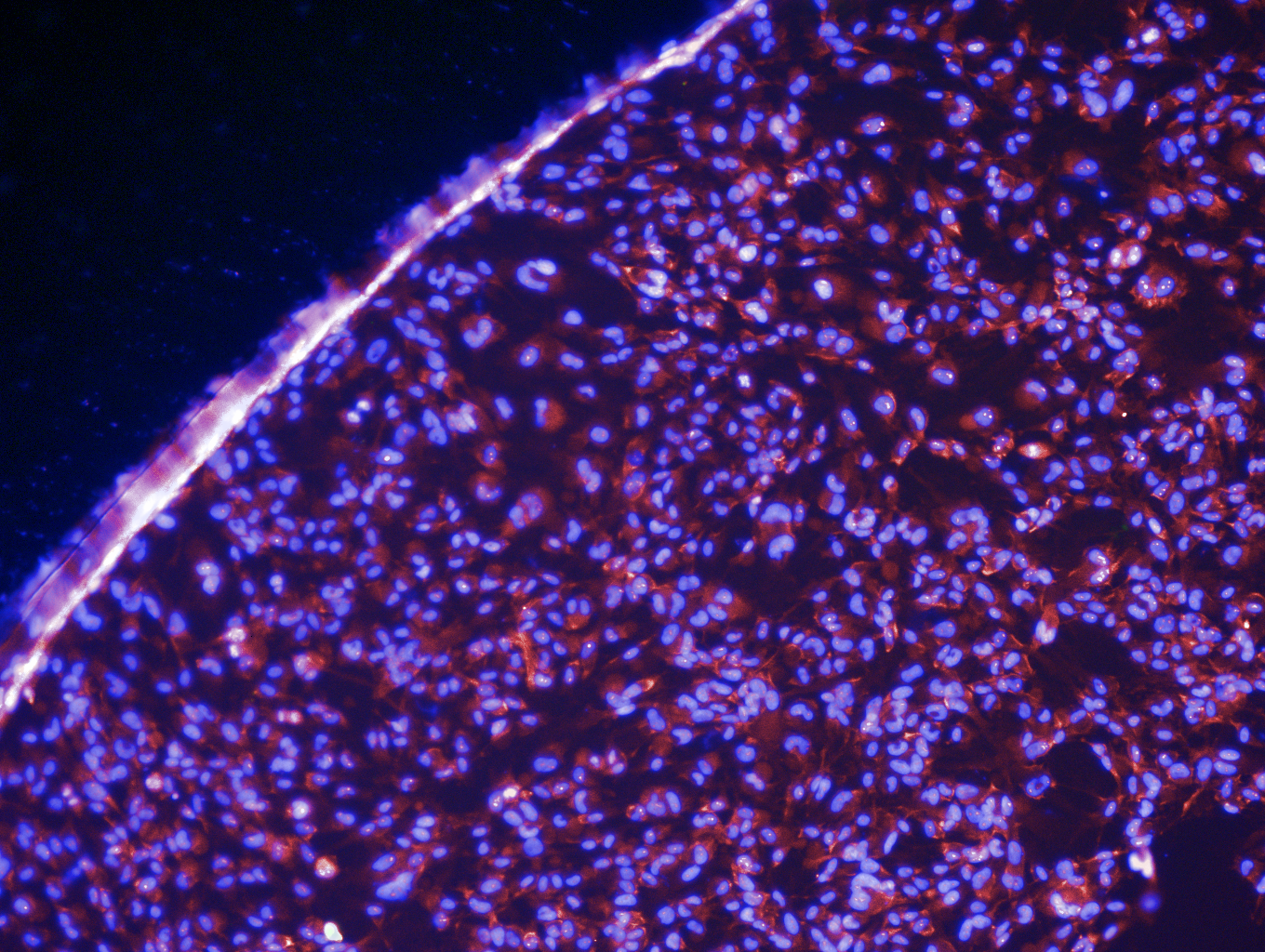

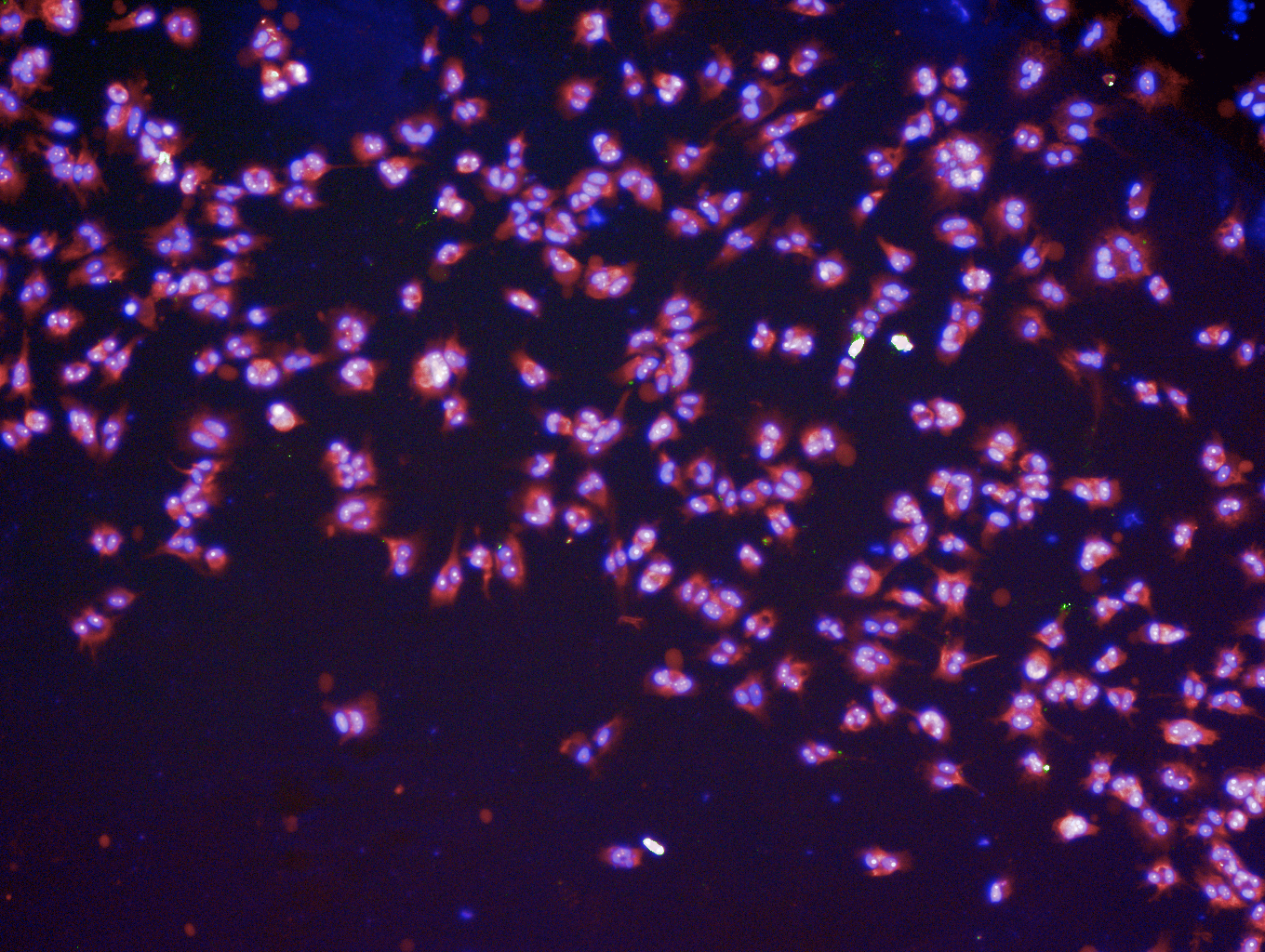

Negative control

miR-1300

U251

Negative control

miR-1300

KNS42

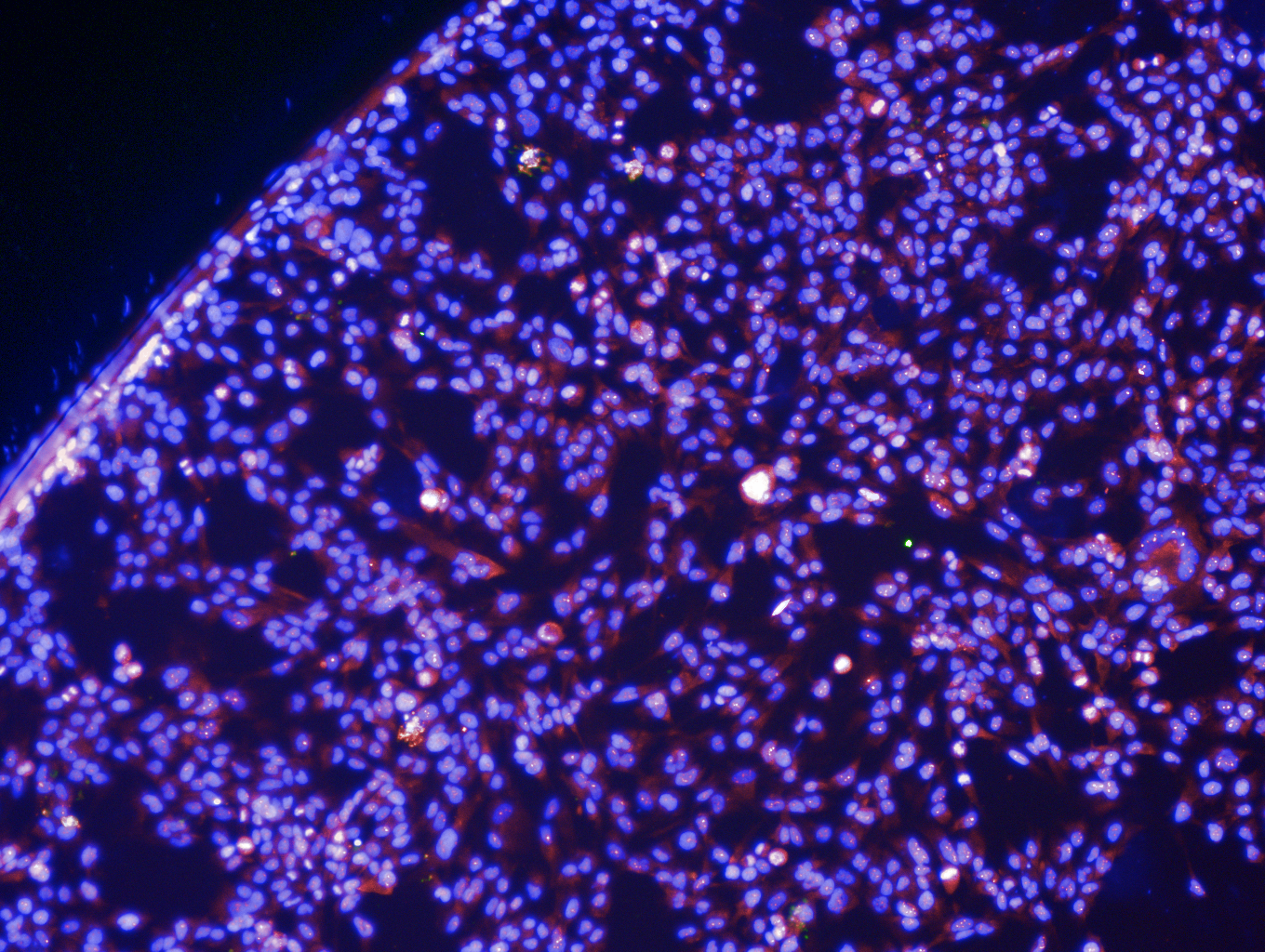

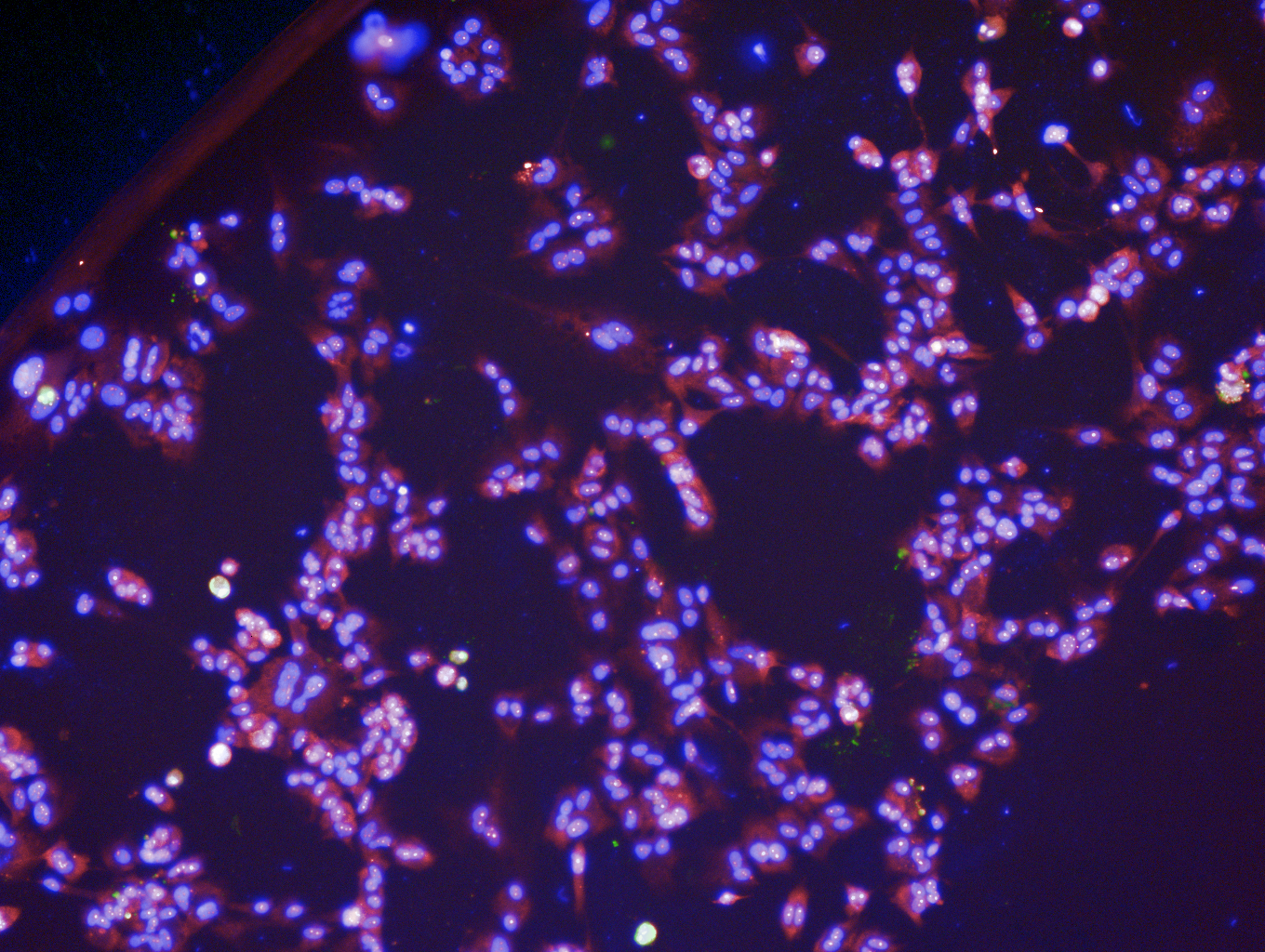

Supplementary Figure 2 **a**: Confirmation of the cytotoxic effect of all seven initial microRNA mimics chosen for validation in a panel of three adult and one pediatric glioblastoma cell line (U251, LN229, U373 and KNS42 respectively). Results are expressed as z-scores calculated as described in the methods section dedicated to the high-throughput screen. **b and c**: Effect of all seven candidate hit microRNA mimics on programmed cell death. Apoptosis was measured by counting Cleaved-Caspase 3 foci by immunocytochemistry, Operetta imaging and custom Columbus software analysis. Cells were stained with DAPI (blue; nuclei) and TOTO-3 (red, cytoplasm). Scale bar =100m. Results clearly show that from the seven candidate microRNAs, miR-1300 is the sole microRNA to induce apoptosis in all cells tested. All experiments were performed in three times and contained triplicate wells for each condition (Untreated, Negative control, or microRNA mimic).

**MicroRNA-1300 precursor sequence**

**5’-**CACCACUGCUGGCCAUCUGAUCUACAAAUGCAGUGGCAUUGACAAAAGAAC

CAUUGAAAAU**UUGAGAAGGAGGCUGCUG**AGAUGGGA-3’

**MicroRNA-1300 mature sequence and seed region**

U**UGAGAAG**GAGGCUGCUG

**Blast against genomic database:** perfect **Plus/Plus strand** homology to EEF1A1P22, pseudogene of EEF1A1- chr 15, thus indicating that the precursor might be expressed from the “minus strand of pseudogene22” on chr15.

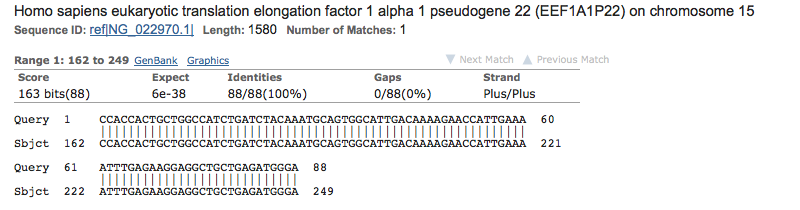

**Ensembl page for EEF1A1P22**: known processed pseudogene from which miR-1300 is expressed.

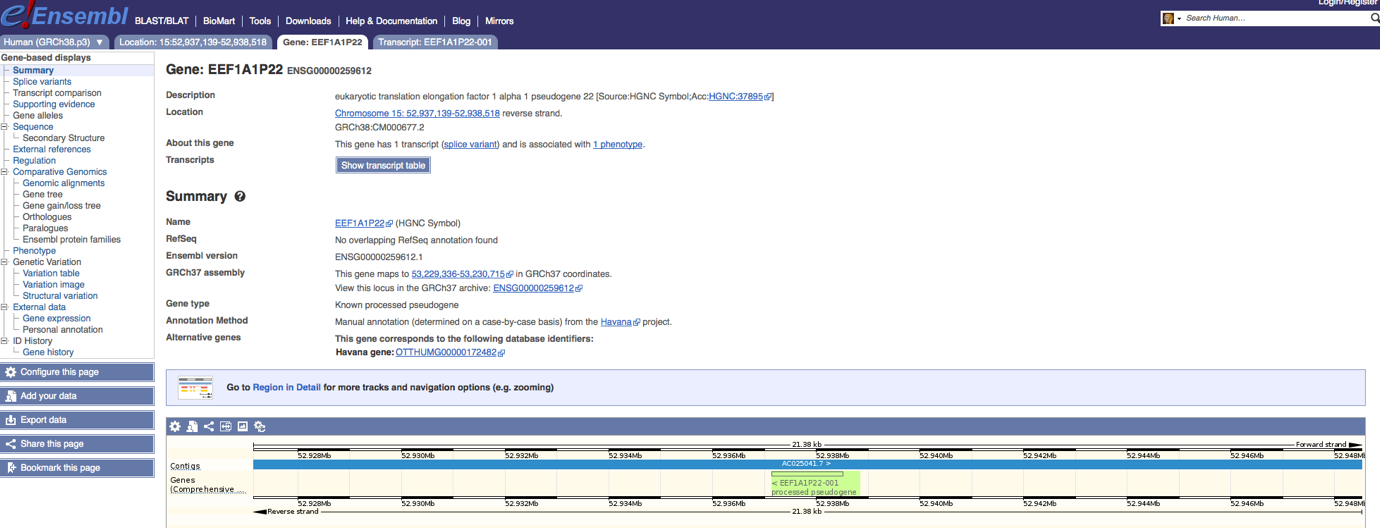

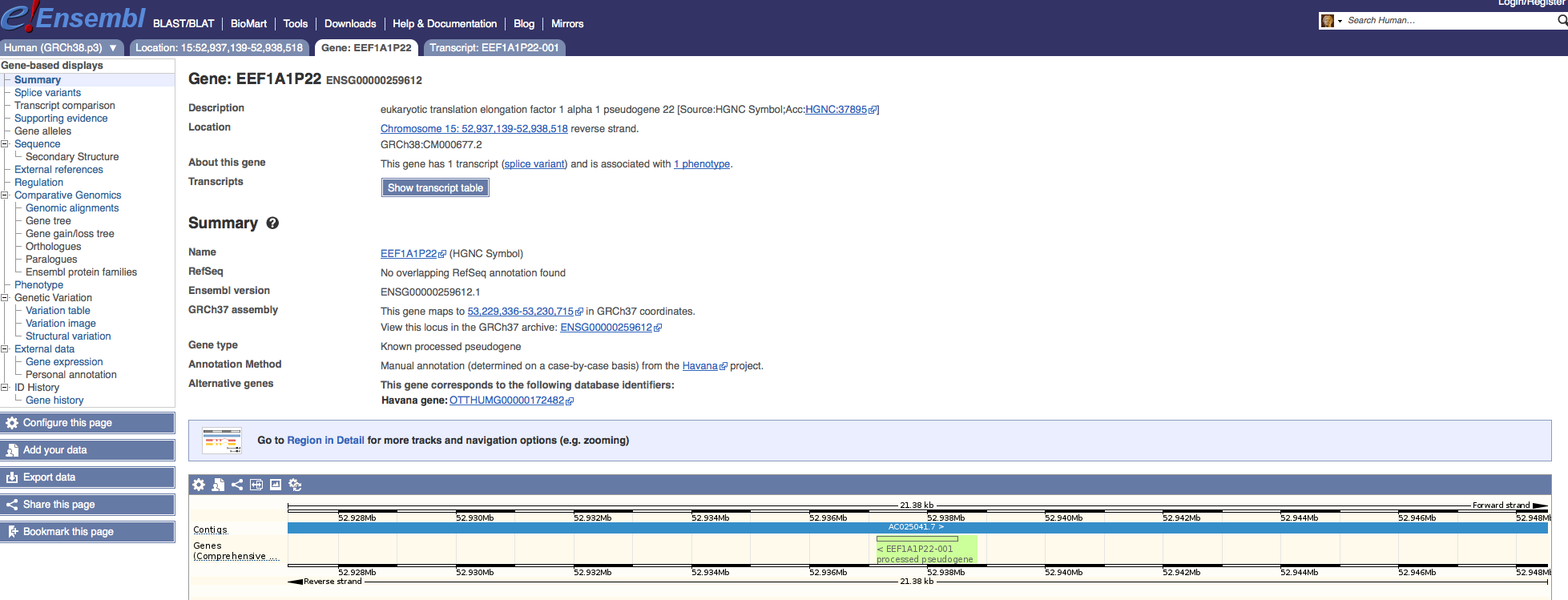

Supplementary Table 4: MiR-1300 sequences and alignment to the human genome.

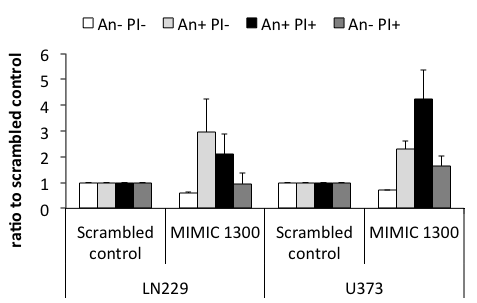

**

*

*

***

**

****

*

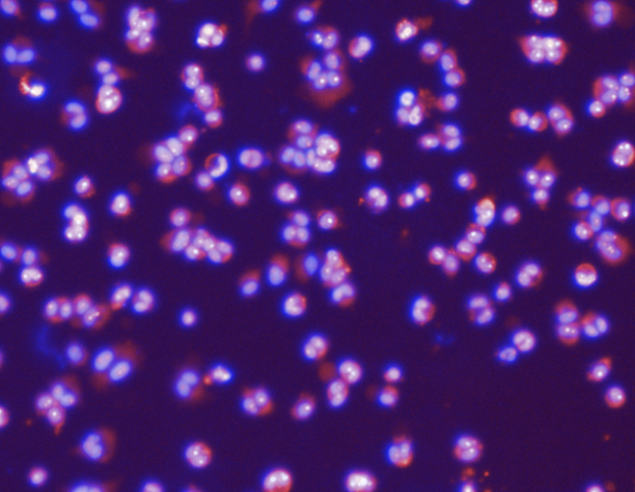

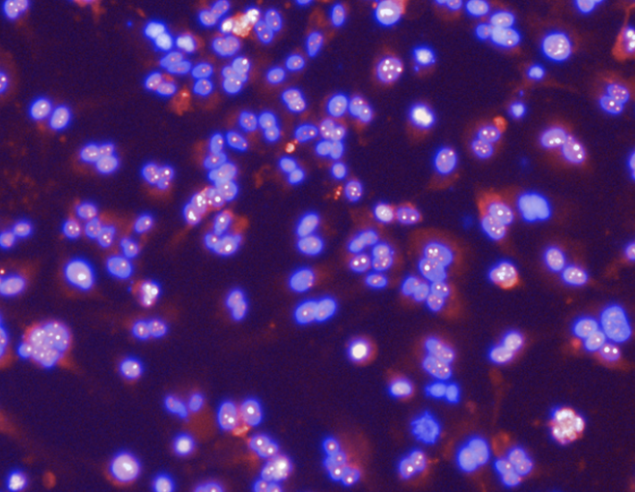

**LN229**

**U373**

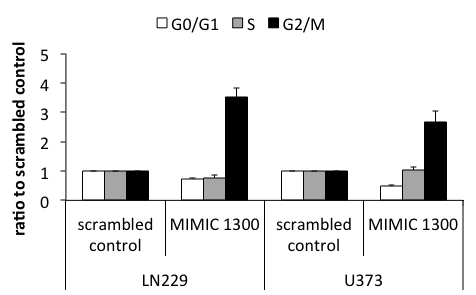

**

***

****

***

**a**

**c**

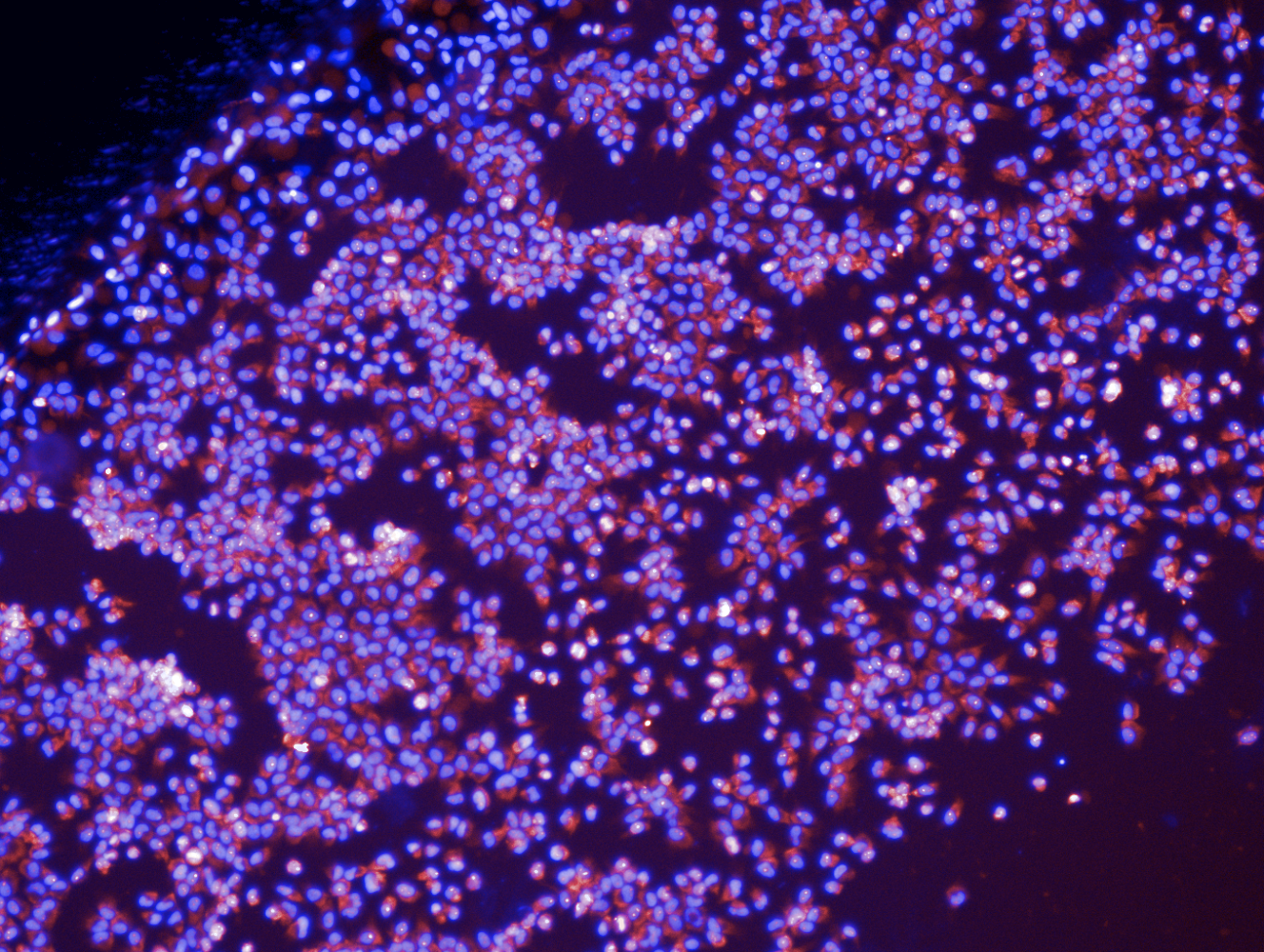

Scrambled control

MIMIC miR-1300

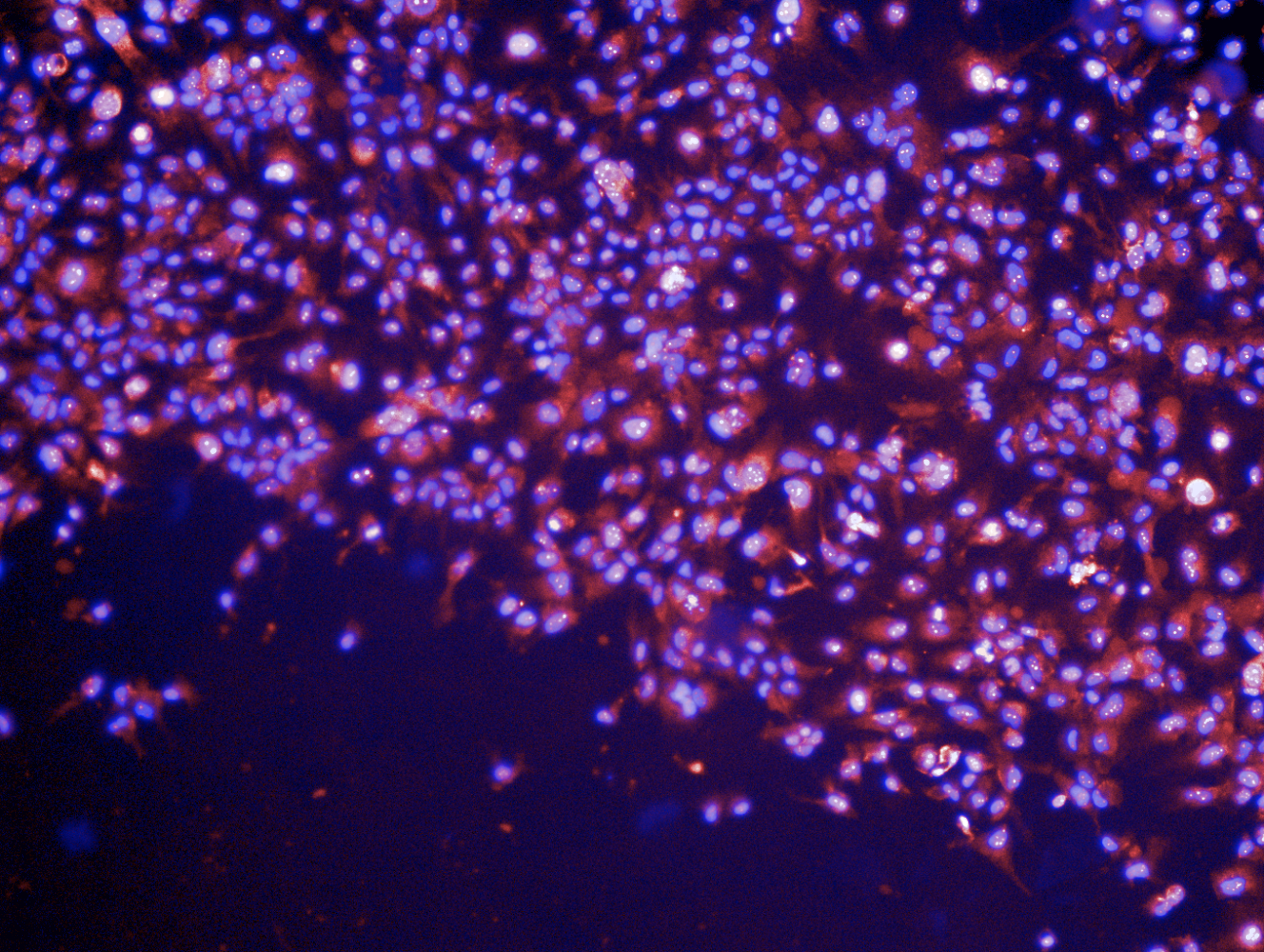

Scrambled control

MIMIC miR-1300

**b**

**100µm**

**100µm**

**100µm**

**100µm**

Supplementary Figure 3: Ectopic expression of miR-1300 causes cell cycle arrest, apoptosis and binucleation in LN229 and U373 adult GBM cell lines 72h post-transfection.

**a**: Effects on the cell cycle were analyzed by flow cytometry using Propidium Iodide (PI) loading as a measure of DNA content. **b**: Cell death measured by flow cytometry where Annexin negative (An -) and PI negative (PI-) = live cells, An+ PI- = early apoptotic cells, An+ PI+ = mid/late apoptotic cells and An- PI+ = Necrotic cells. **c**: Binucleation phenotype scoring (3-6 images per condition, representing at least 100 cells) was performed on images taken on the Operetta imaging platform (x10 objective). Here, representative images show cells stained with DAPI (DNA dye, marking the nucleus) and TOTO-3 (RNA dye, showing the extent of the cytoplasm) All experiments were performed in triplicate.. Statistical significance is expressed as follows: * = p<0.05, ** = p<0.01, *** = p<0.001 and **** = p<0.0001. Scale bar =10m.

**Supplementary movies links and legend**

<http://www.bioinformatics.leeds.ac.uk/~bs06lw/BoissinotEtAl_Videos/AdditionalMovie1A.mp4>

Supplementary Movie 1A (Boissinot et al. Suppl. Movie 1A.mp4) Live imaging of KNS42 cells taken on a Nikon Biostation IM using a x20 objective showing the effect of 100nM miR-1300. Observe a failure in abscission after telophase. Cytokinesis failure becomes increasingly apparent with time post-transfection.

<http://www.bioinformatics.leeds.ac.uk/~bs06lw/BoissinotEtAl_Videos/AdditionalMovie1B>[.mp4](http://www.bioinformatics.leeds.ac.uk/~bs06lw/BoissinotEtAl_Videos/AdditionalMovie1A.mp4)

Supplementary Movie 1B (Boissinot et al. Suppl. Movie 1B.mp4) Live imaging of U251 cells taken on a Nikon Biostation IM using a x20 showing the effect of 100nM miR-1300. Observe a failure in abscission after telophase. Cytokinesis failure becomes increasingly apparent with time post-transfection.

ARHGEF11

PRC1

DIAPH2

IGF1R

RALA

DRD2

ARL3

CETN2

CALM3

CALM1

CUL3

RAB35

PKN2

RAB11FIP4

FLCN

CKAP2

E2F7

NUSAP1

ECT2

BIN3

FMN2

Supplementary Table 5 (Boissinot at al. Suppl. Table 3.pdf) List of miR-1300 predicted target genes associated with cytokinesis pathways.

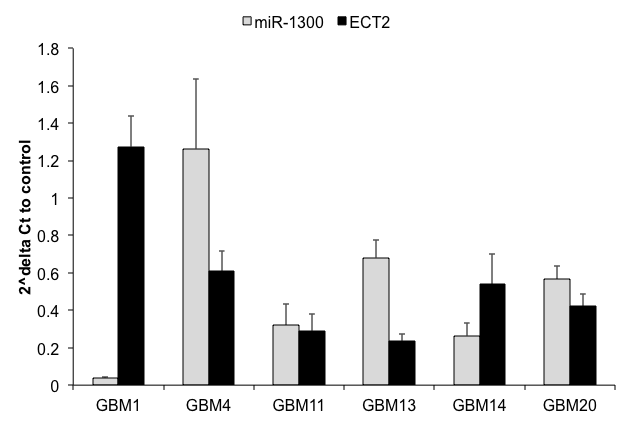

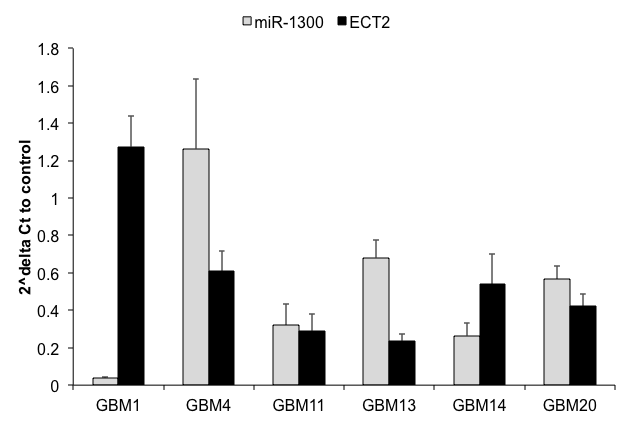

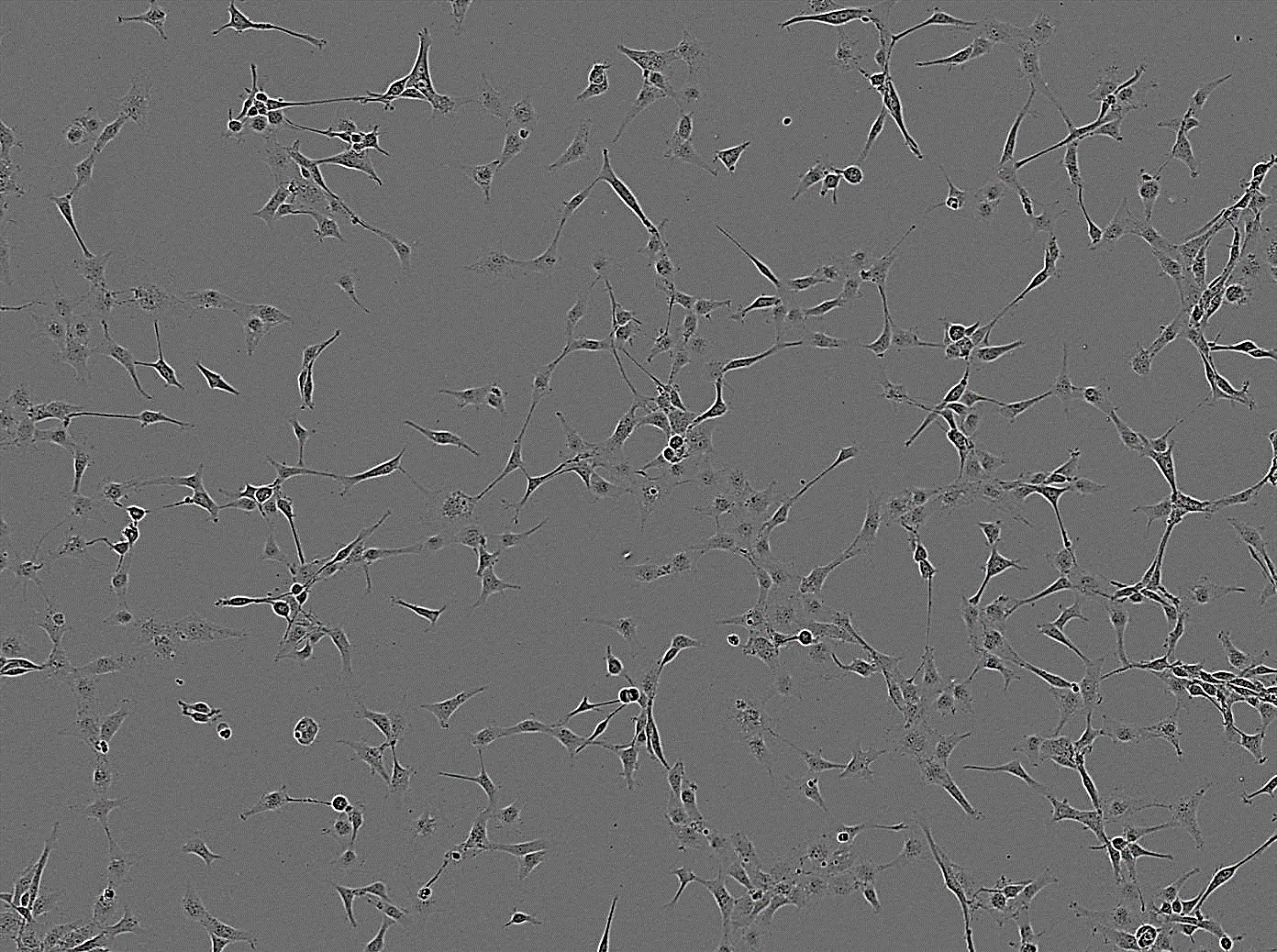

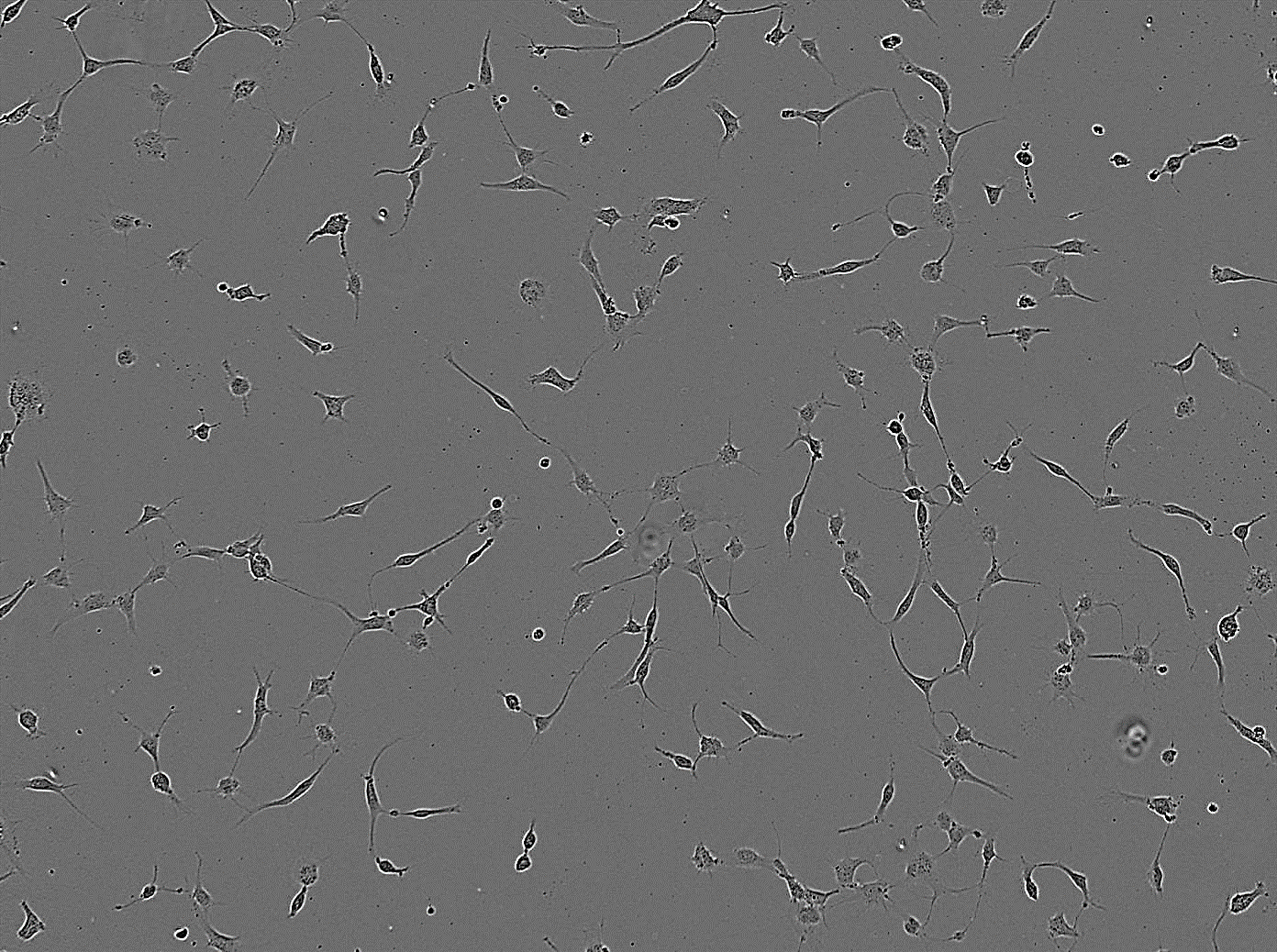

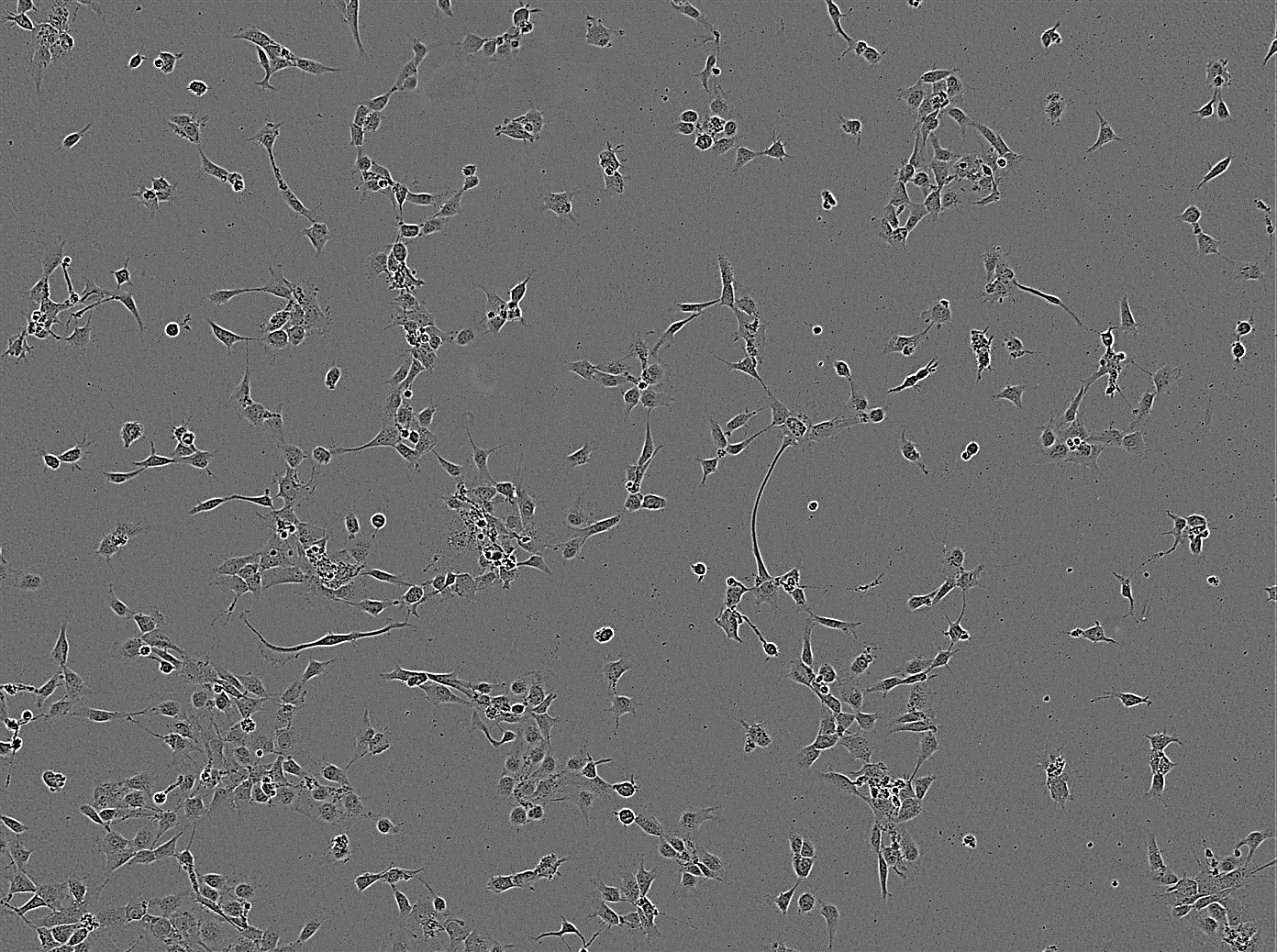

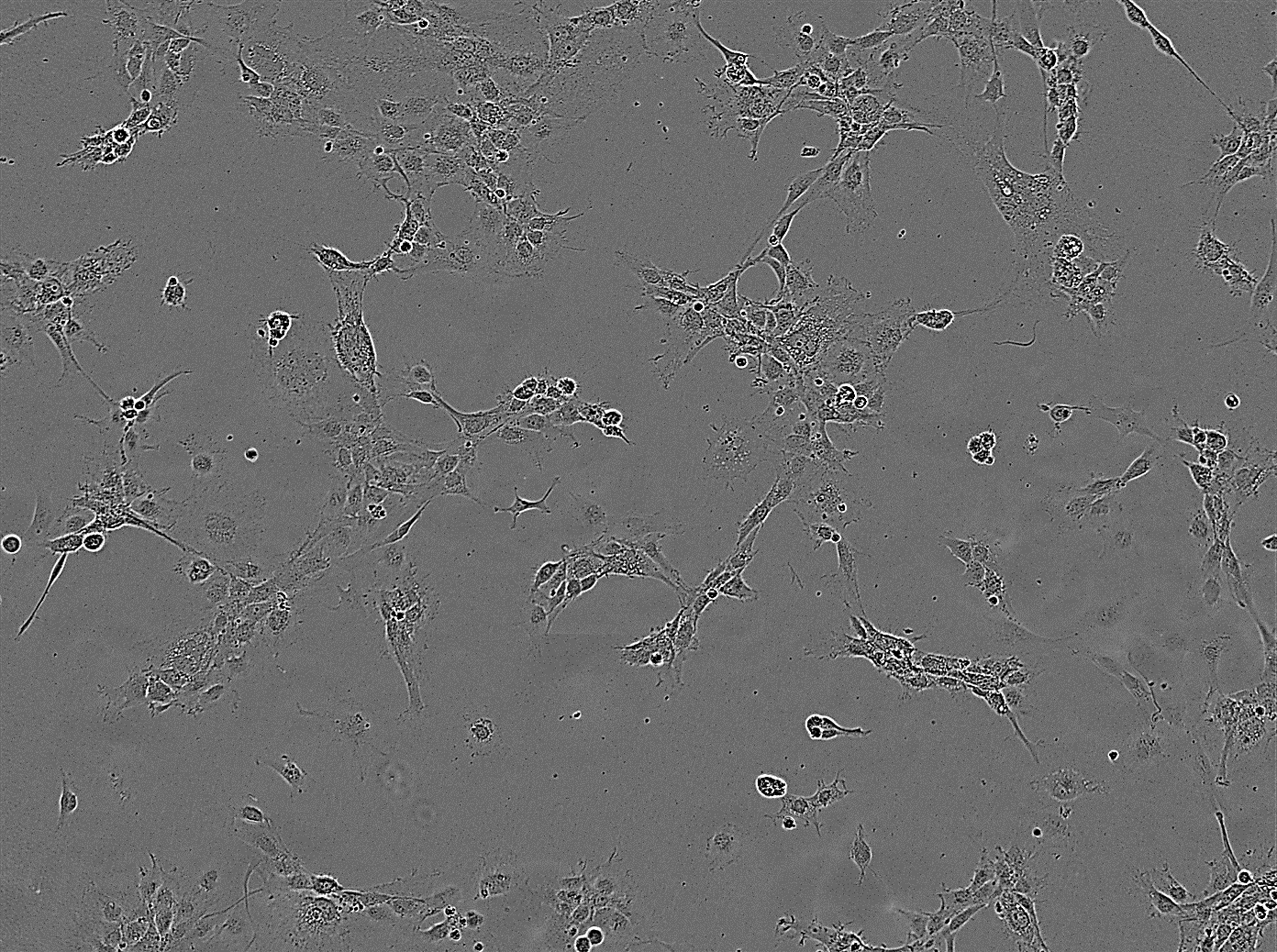

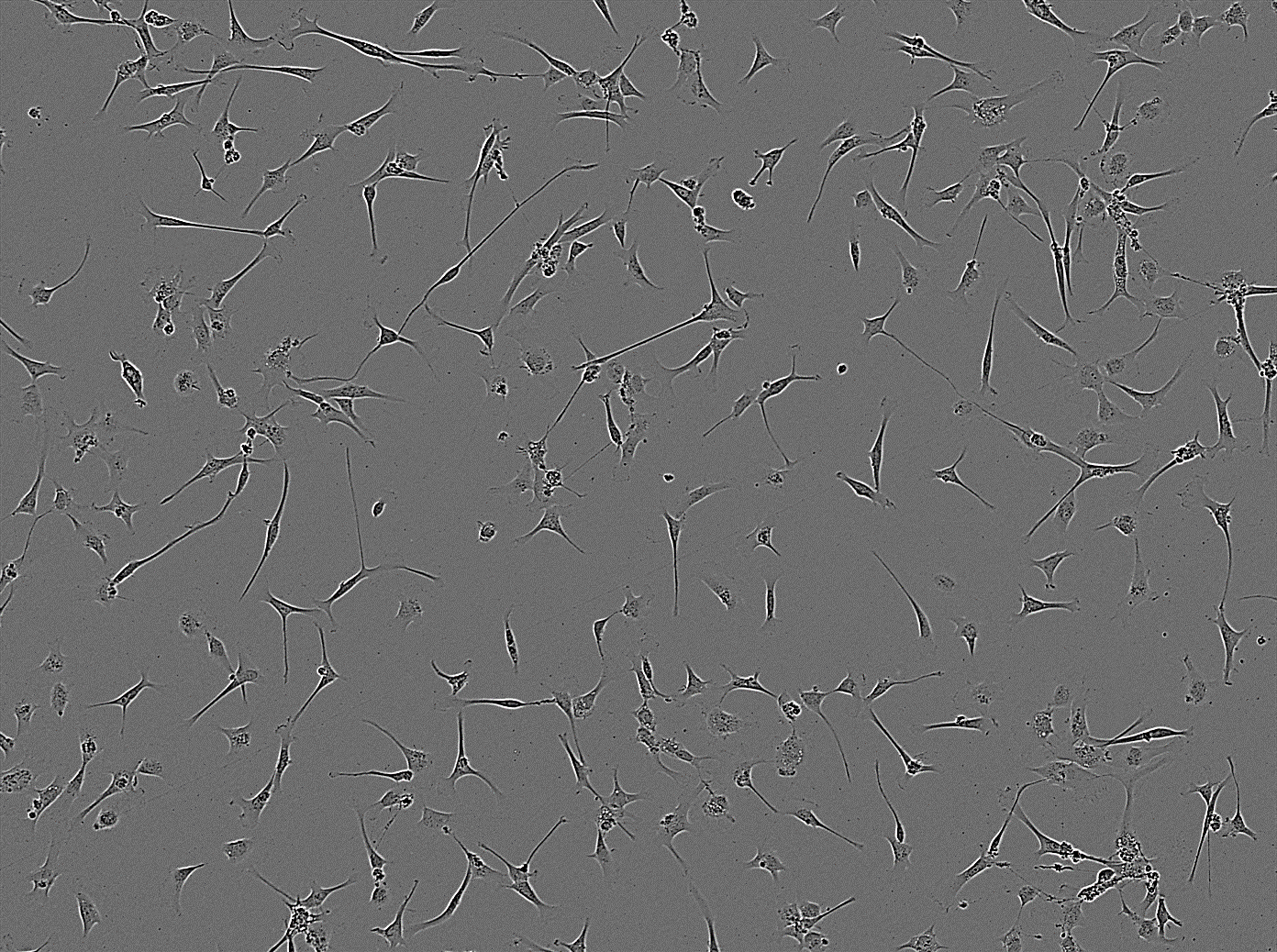

**GBM1	GBM4	GBM11**

100 µm

**GBM13	GBM20**

100 µm

100 µm

100 µm

100 µm

Supplementary Figure 4: A. Endogenous levels of mature miR-1300 and ECT2 mRNA expression measured by Real Time qPCR in a panel of stem-like patient-derived GBM cell lines. ECT2 and miR-1300 expression tends to be inversely proportionate to each other. This also corresponds to frequency of multinuclear cells observed in culture (B).

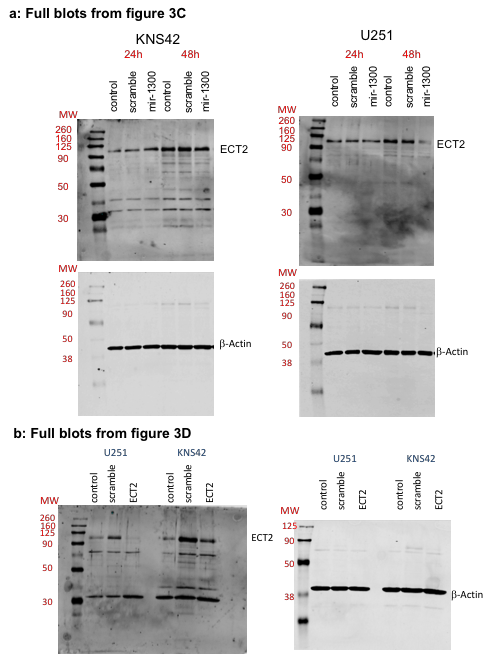

Supplementary Figure 5 **a**: Full blot images for ECT2 and β-Actin corresponding to figure 3C. **b**: Full blot images for ECT2 and β-Actin corresponding to figure 3D. Whole cell lysates were prepared and separated by SDS-PAGE, electroblotted to a nitrocellulose membrane (BioRad) and probed with anti-ECT2 (Santa Cruz, Cat #sc-1005) used at 1/200 dilution. Signal was detected using the Supersignal West Pico Rabbit IgG detection kit at 1/5000 (Cat# 34083, Thermo Scientific). Antibody binding was detected using the Odyssey imaging system (LI-COR Biosciences). Equal lane loading was confirmed using a monoclonal antibody against β-actin (Sigma-Aldrich, Cat#A1978) used at 1/10,000.

All experiments were performed in triplicate. Results were normalized to a scrambled mimic control. Statistical significance is expressed as follows: ** = p<0.01, **** = p<0.0001. NB: Blot images were taken at 48h time point since at 72h, cells in control conditions have reached confluence, and were not expressing ECT2 anymore as they are not dividing.

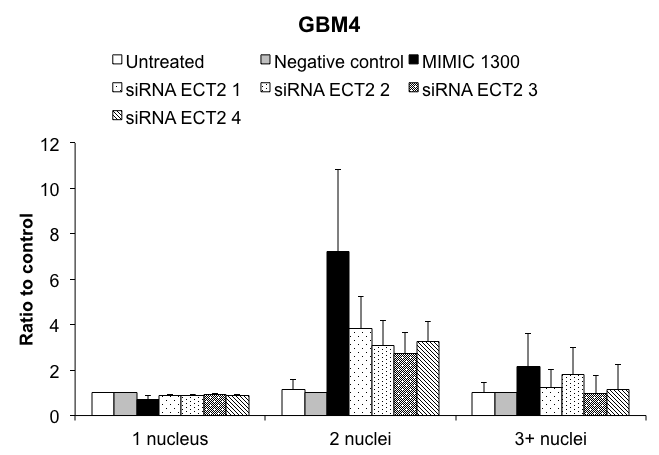

****

*

****

****

****

****

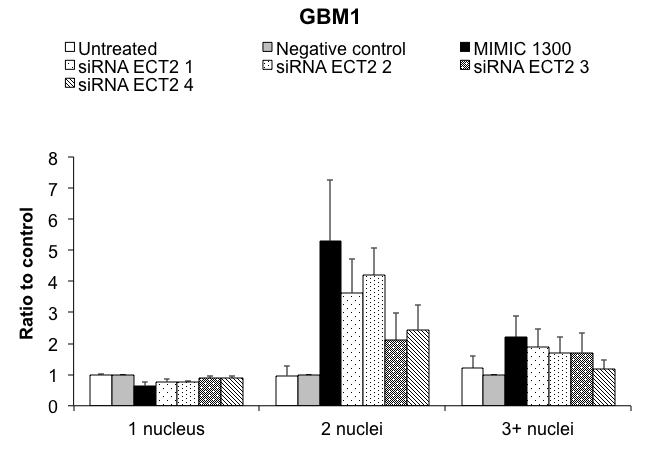

****

**

***

**

**

*

*

****

****

Supplementary Figure 6: ECT2 knock-down in stem-like GBM1 and GBM4 cells. Individual siRNAs against ECT2 were transfected in patient-derived GBM1 and GBM4 cells. Immunofluorescence and binucleation scoring were performed at 96h post-transfection. 3 images per replicate per condition were scored, and experiments performed in triplicates. Results are normalized to a scrambled mimic control. Statistical significance is expressed as follows: * = p<0.05, ** = p<0.01, *** = p<0.001 and **** = p<0.0001.

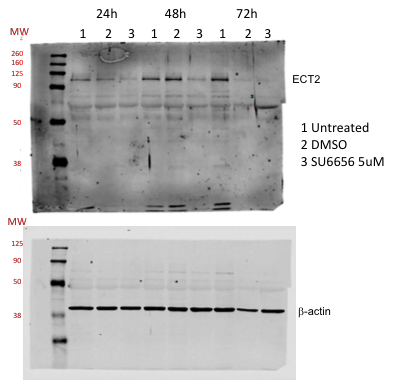

Supplementary figure 7: Full blot images for ECT2 and β-Actin corresponding to figure 6. Exposure of synchronised CMK cells to 5uM of SU6656 concomitantly induces an increase in the levels of miR-1300 and a decrease of its targets, ECT2. Whole cell lysates were prepared and separated by SDS-PAGE, electroblotted to a nitrocellulose membrane (BioRad) and probed with anti-ECT2 (Santa Cruz, Cat #sc-1005) used at 1/200 dilution. Signal was detected using the Supersignal West Pico Rabbit IgG detection kit at 1/5000 (Cat# 34083, Thermo Scientific). Antibody binding was detected using the Odyssey imaging system (LI-COR Biosciences). Equal lane loading was confirmed using a monoclonal antibody against β-actin (Sigma-Aldrich, Cat#A1978) used at 1/10,000.

All experiments were performed in triplicate.

Supplementary methods:

**Cell lines**:

Established lines: U251, U87 and U373 cell lines were purchased from the Health Protection Agency Culture Collections (Cat# 09063001, 89081402 and 08061901 respectively). LN229 cells were purchased from the American Type Culture Collection (Cat# ATCC-CRL-2611). KNS42 cells were a gift from Prof. C. Jones (Institute of Cancer Research). Cells were mycoplasma free and authenticated in house by STR profiling. Cells were grown as monolayers in DMEM 4.5mg glucose, Glutamax (Sigma D6429), 10% heat-inactivated FCS, 1X penicillin/streptomycin.

Patient-derived GBM stem cells: GBM1 and GBM4 were isolated from patient tumor material and maintained as adherent cultures on poly-L-ornithine/laminin coated substrate in Neurobasal medium (Life Technologies) supplemented with; N2 and B27 (0.5 x each, Life Technologies) and recombinant bFGF and EGF (40 ng/ml, R & D systems) as previously described (23-25).

The CMK megakaryocytic cell line CMK (CSC-C0439) was purchased from Creative Bioarray (New York, USA) and cultured in RPMI-1640, 20% heat-inactivated FCS, 1X penicillin/streptomycin. Prior to experiments, cells were synchronised using Monastrol at 25μM for 24 hours.

**Bioinformatics analysis:**

The list of miR-1300 predicted targets (from TargetScan v5.2) was analyzed using Metacore (Thomson Reuters) and AmiGO (28) to match “cytokinesis” as a Gene Ontology (GO) term with our gene list. The results were further analyzed in a step wise manner allowing for the exclusion of genes coding for non-functional proteins. Finally, we intersected these data with the initial list from TargetScan. All analysis was performed using “R”.

**qRTPCR:**

MicroRNAs were extracted from cells using the miRNeasy Mini kit (Cat# 217004, Qiagen) following manufacturer’s instructions. The small RNA enrichment step was used for experiments with microRNA/mRNA co-extraction.

MicroRNA reverse transcription was performed using the specific ABI kit (Cat# 4366596) and the matched reverse transcription and TaqMan primer sets for each microRNA and the housekeeping control U6snRNA were supplied by Applied biosystem (ABI) by Life Technologies Cat#442975 then assay # 002902 for hsa-miR-1300 and assay # 001973 for U6snRNA, respectively.

When only mRNAs were required, extraction was performed using a classic non-column based Trizol/Chloroform/precipitation method. Reverse transcription was then performed using the Superscript II reverse transcriptase (Cat#18064014), oligodT primers (Cat#18418-012), dNTPs (Cat# 18427-013) and RNAse OUT (Cat# 10777019) from Life Technologies.

Predesigned assays including primers and 5' 6-FAM / 3' TAMRA probe were purchased from Integrated DNA Technologies: ECT2 (**Hs.PT.58.38680333),** HPRT1 (housekeeping) (Hs.PT.58v.45621572).

All RTqPCR mixes were prepared using the TaqMan PCR MixII no UNG (Cat# 444048) in a 96 well plate format (Cat#4306737) and run on a 7500HT thermocycler all from Applied Biosystems.

**Immunofluorescence staining for candidate validation:**

72h post transfection, cells were washed and fixed in 4% paraformaldehyde for 20 mins at room temperature for Phalloidin (actin) and cleaved-caspase-3 staining or with ice-cold methanol for α-tubulin staining. Cells were then washed in PBS and permeabilized for 3 mins in PBS containing 0.1% Triton-X-100. Non-specific binding was abrogated by a blocking step of 5 mins in 1% BSA in PBS. Cells were then stained for 1h at room temperature with either: 5μM of Phalloidin Alexafluor488 (Cat♯ A12379, Life Technologies) or 1/3000 primary rabbit anti-cleaved Caspase-3 antibody (Cat♯ 700182, Life Technologies) or 1/500 rat anti-α-tubulin (Cat♯ MCA77G, AbDSerotec). After three washes in PBS, the following secondary antibodies conjugated with Alexafluor488 and DAPI at 1/4000 (Cat. D1306, Life Technologies) were added for 1h at room temperature: for cleaved-Caspase-3, 1/1000 goat anti-rabbit; for α-tubulin, 1/500 donkey anti-rat. This step was followed by 3 x 5 minutes washes in PBS. Cells were then kept in PBS at 4°C until imaging. Manual scoring of the binucleation phenotype (3-6 images per condition, representing at least 100 cells) was performed on images from the control and candidate hit wells taken on the Operetta imaging platform using 10x objective.

**Immunofluorescence, image capture and high content image analysis for the assessment of endomitotic CMK**

To fix, stain and analyze SU6656 treated non-adherent CMK cells, 2ml of cells in media were transferred to a 15ml falcon tube and fixed by incubation for 30mins with 2ml of 4%PFA with gentle agitation. For each centrifugation step, cell suspensions were centrifuged at 1000g for 30 seconds and the supernatant removed using a pipette. After spinning down the cells, the pellet was gently washed in 2ml PBS, centrifuged and the pellet resuspended in 1ml PBS-0.1% Triton X-100 for 5mins at room temperature. Pelleted cells were washed in 1ml PBS, centrifuged and the supernatant removed before the addition of 1ml of PBS-1% powdered milk blocking solution and incubation at room temperature for 5mins. Cells were pelleted and incubated for 1 hour in the dark in 200l staining solution containing 1:1000 DAPI (Cat♯ D1306, Life Technologies) and 1:2000 (Cat♯ T-3604, Molecular Probes) TOTO3 in 1% powered milk in PBS, prior to washing cells three times in PBS. Cells were pelleted and resuspended in 150l PBS before transferring to individual wells of a 96 well ViewPlate (Perkin Elmer). Cells were allowed to settle to the bottom of the plates before being imaged on the Operetta using Harmony 3.1 software, 2 fluorescent channels, 9 fields/well with a 20x objective. The experiment was performed for at least 3 biological repeats.

Algorithms for the detection of cell number and for polyploidy were created using Columbus 2.4 analysis software. Polyploidy was detected using the cell identification parameters of (i) Find cells (TOTO3 channel), (ii) Select population (remove border objects to create a whole cell population, (iii) Calculate area and roundness morphology properties in whole cell population, (iv) Filter whole cell population based on cell area and roundness to remove platelet populations, (v) Select the megakaryocyte (MKC) cell population based on cell area and roundness, (vi) Select the megakaryocyte (MKB) cell population based on cell area and roundness.

**Flow cytometry assays:**

*Cell cycle PI:*

Briefly, cells were fixed and permeabilized in freezing cold PBS 70% ethanol, washed and then stained by a solution of RNAseA/Propidium Iodide and analyzed on the Attune Flow cytometer (ThermoFisher). Analysis was performed using the specific algorithms in Modfit LT 3.2 (Verity software for ploidy and cell cycle analysis).

*AnnexinV/PI:*

Becton Dickinson assay number: Apoptosis Kit I (Cat# 556547), was used according to manufacturer’s instructions. Cells were then run on the Attune Flow cytometer. Analysis was performed using the Attune software.

**siRNA knock-down:**

Reverse transfections were performed as described above (see § “high-throughput screen) using either ON-TARGET plus *ECT2* SMARTpool (Cat# L-006450-00-0005) which encompasses 4 different siRNA sequences in a single well or each of the four separate siRNA sequences (supplementary figure 6) (Dharmacon from GE Healthcare). Post incubation, cells were fixed and stained as described above.

**Western Blotting:**

siRNA knock-down was confirmed by Western blotting. Whole cell lysates were prepared and separated by SDS-PAGE, electroblotted to a nitrocellulose membrane (BioRad) and probed with anti-ECT2 (Santa Cruz, Cat #sc-1005) used at 1/200 dilution. Signal was detected using the Supersignal West Pico Rabbit IgG detection kit at 1/5000 (Cat# 34083, Thermo Scientific). Antibody binding was detected using the Odyssey imaging system (LI-COR Biosciences). Equal lane loading was confirmed using a monoclonal antibody against β-actin (Sigma-Aldrich, Cat#A1978) used at 1/10,000.
